# Parallel but distinct adaptive routes in the budding and fission yeasts after 10,000 generations of experimental evolution

**DOI:** 10.1101/2025.09.11.675703

**Authors:** Arnaud N’Guessan, Vivian Wang, Christopher W. Bakerlee, Jenya Belousova, Greta Brenna, Megan E. Dillingham, Thomas Dupic, Shreyas Gopalakrishnan, Juhee Goyal, Misha Gupta, Caroline Holmes, Parris T. Humphrey, Tanush Jagdish, Elizabeth R. Jerison, Milo S. Johnson, Katya Kosheleva, Katherine R. Lawrence, Jiseon Min, Alief Moulana, Shreyas V. Pai, Angela M. Phillips, Julia C. Piper, Ramya Purkanti, Artur Rego-Costa, Tatiana Ruiz-Bedoya, Cecilia Trivellin, Michael J. McDonald, Michael M. Desai, Alex N. Nguyen Ba

## Abstract

Quantitative genetics approaches designed to understand the evolution of traits have helped improve our understanding of the genetic basis of adaptation. However, they often overlook crucial aspects of adaptation, including the long-term temporal evolutionary dynamics, the predictability of evolutionary outcomes, the influence of past evolution on future evolutionary trajectories (contingency), and the diversity of molecular mechanisms underlying adaptation. Experimental evolution has been a useful tool for answering these questions, but extracting fundamental principles and predictive features of evolutionary outcomes from these datasets remains challenging due to the large number of covariates and confounding effects, such as differences in experimental setup, species lifestyle, gene content, and evolution rate. Here, we sought to circumvent these challenges by comparing distant yeast species that share several evolutionary features but differ mainly in evolutionary history and genome architecture, i.e. *Saccharomyces cerevisiae* and *Schizosaccharomyces pombe*. Thus, we evolved 10 populations of the fission yeast for 10,000 generations in the same conditions as a pre-existing budding yeast dataset (i.e. high-sugar media and hypoxic conditions), allowing us to observe repeatable evolutionary outcomes within species but diverse molecular mechanisms and targets of adaptation across species. The most frequent adapting route in these conditions involved upregulating fermentation genes and downregulating the glycolysis gene *pyk1*, which has not previously been observed in *S. cerevisiae* evolved populations or in wild *Kluyveromyces lactis,* but similar evolutionary paths have been observed in *Schizosaccharomyces japonicus* and in clinically relevant populations, such as some cancer cells. This suggests that parallelism is pervasive in the tree of life and that mechanisms of adaptation can be shared among closely related or distant species. Despite similar gene content and identical environments, recurrent adaptation across *S. pombe* populations involved different genes than in *S. cerevisiae* and was mostly detectable at the transcriptomic level. This suggests that trans-regulatory effects may play an important role in adaptation on short evolutionary timescales and that differences in evolutionary outcomes between these species may be attributed to contingency.

## INTRODUCTION

Long-term experimental evolution has highlighted the predictability of high-level features of evolution like fitness trajectories^1–6^. While fitness gains follow a fairly reproducible pattern of declining adaptability, the exact mutations or the sequence in which they occur are typically unpredictable. Nevertheless, experimental evolution done in large replication with the budding yeast *Saccharomyces cerevisiae* has revealed extensive parallelism at the gene or pathway level^3,7,8^, even across strains that differed by thousands of polymorphisms^9^. These general patterns of reproducibility during experimental evolution are also seen in different organisms. For instance, during the *Escherichia coli* Long-Term Experimental Evolution (LTEE), half of the populations evolved a mutator phenotype and quasi-stable lineage coexistence emerged in 75% of the populations^4^. Further, substantial parallelism at the gene level can also be observed throughout the 12 populations over (now greater than) 60,000 generations^4^.

Unsurprisingly, the molecular targets of selection during these experiments differ substantially between *S. cerevisiae* and *E. coli*^3,6^. Likewise, comparing microbial laboratory evolution to experimental evolution done in *Caenorhabditis elegans* and *Drosophila melanogaster* does not reveal universal targets of adaptation^10,11^. Notwithstanding their vastly different gene content and cellular form, part of the reason for these differences in molecular adaptation is the different selection environments between experiments (the LTEE was done in DM25 minimal media, while yeast was done in YPD rich media or in synthetic dextrose complete media). However, another more interesting hypothesis for these differences relates to the role of evolutionary contingency in shaping the dynamics and targets of selection.

Contingency is the product of past evolution, whereby epistasis and genome structures can strongly influence future adaptive potential^1,12,13^. Thus, species can be constrained to alternative adaptive paths, leading to profoundly different molecular signatures of evolution. Contingency has been explored in the context of strains with different gene deletions^13–16^ and has even been observed in experimental evolution. For instance, across the 12 *E. coli* populations evolving in the same conditions during the LTEE, one population developed the ability to grow aerobically on citrate, a rare phenotype in this species resulting from a rare genome rearrangement. This phenotype is far more readily evolvable within that population after this rearrangement^17^. Further, while half of the LTEE populations evolved a mutator phenotype, and 75% evolved stable lineage coexistence, these phenomena are not observed in *S. cerevisiae* laboratory evolution experiments in standard rich media or in minimal media, despite being entirely capable of achieving these through genetic engineering^18^.

While studying evolution using very distinct organisms has provided important insights into general features of evolutionary dynamics, it has made it difficult to discern the effect of life history from other idiosyncrasies on the molecular targets of selection. Would we find the same targets of selection (genes), and a preference for the same molecular mechanisms if we had evolved a species with substantially similar gene contents to that of *E. coli* or *S. cerevisiae*? The difficulty in answering this question is also reinforced by the wide variety of mechanisms of rapid adaptation revealed by long-term evolution studies, e.g. hybridization, selection on standing variation, cross-feeding interactions, or phenotypic plasticity involving behavioral or transcriptomic changes^11^. Evaluating the impact of differences between such species on quantitative trait evolution could improve our understanding of the role of genetic architecture and life history in trait diversity.

One way to address how much these two factors might influence short- and long-term adaptation is through the yeast subphylum. Although it exhibits immense trait diversity and is as genetically diverse as the animal kingdom^19,20^, species within have a comparable number of genes, and share many genes and metabolic pathways such that most species can grow in the same media. As an example, *Saccharomyces cerevisiae* and *Schizosaccharomyces pombe* share about 4,500 genes (out of approximately 6,000), but are so diverged that their genomes show no synteny^21,22^. Evolutionarily, they have similar spontaneous mutation rates and were both domesticated for their ability to ferment. Some notable differences are that *S. pombe* is naturally a haploid in the wild^23,24^, while *S. cerevisiae* is usually found as a diploid^25,26^. This could affect the effectiveness of selection because mutations can be recessive in diploids. Therefore, the extant genome might have ‘footprints’ of this accumulated genetic load in *S. cerevisiae*^3,27^. Supporting this, heterosis has been observed in yeast from crosses of wild budding yeast strains^28,29^. Further, *S. cerevisiae* is within the clade of yeasts that have undergone the so-called ‘whole-genome duplication event’, which is thought to have profoundly shaped their metabolism due to functional divergence in paralogous gene copies^30^. Another major difference is that *S. pombe* cannot survive the loss of its mitochondrial DNA and relies on oxidative phosphorylation for growth, i.e. it is petite-negative, while *S. cerevisiae* can survive without mitochondrial DNA, i.e. it is petite-positive, highlighting a rare ability of budding yeast ^31–33^. Thus, domestication, competition with other microorganisms, and exploration of poorly oxygenated niches might have led *S. cerevisiae* to produce and process ethanol more efficiently than *S. pombe* ^34,35^. Therefore, their different life cycles and evolutionary histories could lead them to different evolutionary outcomes

Hints of the effects of contrasting life and evolutionary histories on evolutionary signatures emerge when comparing the molecular processes that are under selection and could potentially allow for the prediction of molecular evolution that may occur during experimental evolution. For example, *S. cerevisiae* gene content variation across strains is more frequent than in *S. pombe*, and loss-of-function (LOF) mutations, such as nonsense mutations and frameshift deletions, tend to be recurrently fixed across populations^36^. The rate of copy number variants (CNV) is also higher in the budding yeast sub-telomeric regions and can contribute to its adaptation^36^. Additionally, although *S. pombe* has a similar substitution rate to the budding yeast, it has a higher indel mutation rate with a bias toward insertions^37^. Previous genomic analyses also showed that 43% of *S. pombe* genes have introns compared to 5% in *S. cerevisiae* and upstream intergenic regions are larger in the former^38^. The difference in extent of non-coding regions is also present at the whole-genome level: annotated coding regions span around 70% of *S. cerevisiae* genome compared to 60% for *S. pombe*^39–41^. It is currently unknown if this implies that *S. pombe* would preferably adapt through regulatory changes while *S. cerevisiae* would do so through gene content changes and mutations in protein-coding genes. However, the mutations in the non-coding regions of *S. pombe* are known to be involved in transcription regulation and should affect trait evolution and adaptation^42^. Thus, comparing the evolutionary processes and targets of adaptation in *S. cerevisiae* and *S. pombe* through experimental evolution could help highlight factors explaining differences in evolutionary outcomes across and within species.

To address the gap in understanding how species with similar gene content, but different life histories, might evolve in the same environment, we undertook a laboratory evolution experiment where we founded about 16 populations, each of 5 different budding yeast species, as well as the *S. pombe* fission yeast. Here, we describe the dynamics and molecular evolution of the fission yeast populations over the first 10,000 generations. Fission yeast is considered the outgroup of this set of species, and so it serves as a useful anchor for expectations within our chosen fungal phylum. We analyze the mutational spectrum, the molecular evolutionary dynamics, and identify, through pathway enrichment analysis and whole transcriptome sequencing, a case of parallel evolution where *S. pombe* populations have rewired their central carbon metabolism to reduce dependency on oxygen, which mirrors similar events in wild *S. pombe* populations and in related *Schizosaccharomyces*^32,35,35,43,44^.

## RESULTS

### Contrasting fitness gain of fission yeast during 10,000 generations of evolution

To compare the evolutionary processes contributing to adaptation and selection targets between *S. pombe* and *S. cerevisiae*, we founded 15 *S. pombe* populations and performed experimental evolution for 10,000 generations, collecting and freezing samples approximately every 70 generations in 27% glycerol (**Methods; Fig. 1**). The populations were all founded from single colonies of the same *S. pombe* haploid laboratory strain (972h-) and serially propagated in batch culture in one well of a lidded, unshaken flat-bottom 96-well plate in rich media at 30°C with a daily 1:2^10^ dilution. 5 populations were lost during the first 10,000 generations of evolution, due to contamination or mechanical pipetting errors. Because of the plate lidding and the absence of shaking, these conditions correspond to a high-sugar hypoxic environment. The media and temperature are also identical to our previous experimental evolution study, which analyzed 10,000 generations of evolution for *S. cerevisiae* populations that were passaged simultaneously using liquid handling robotics^3^. Although these growing conditions may be sub-optimal for growth of *S. pombe* and so the selection pressure faced by the species may differ, our experiment allows us to control for some of the environmental variate for the comparison of evolutionary outcome.

**Figure 1.**
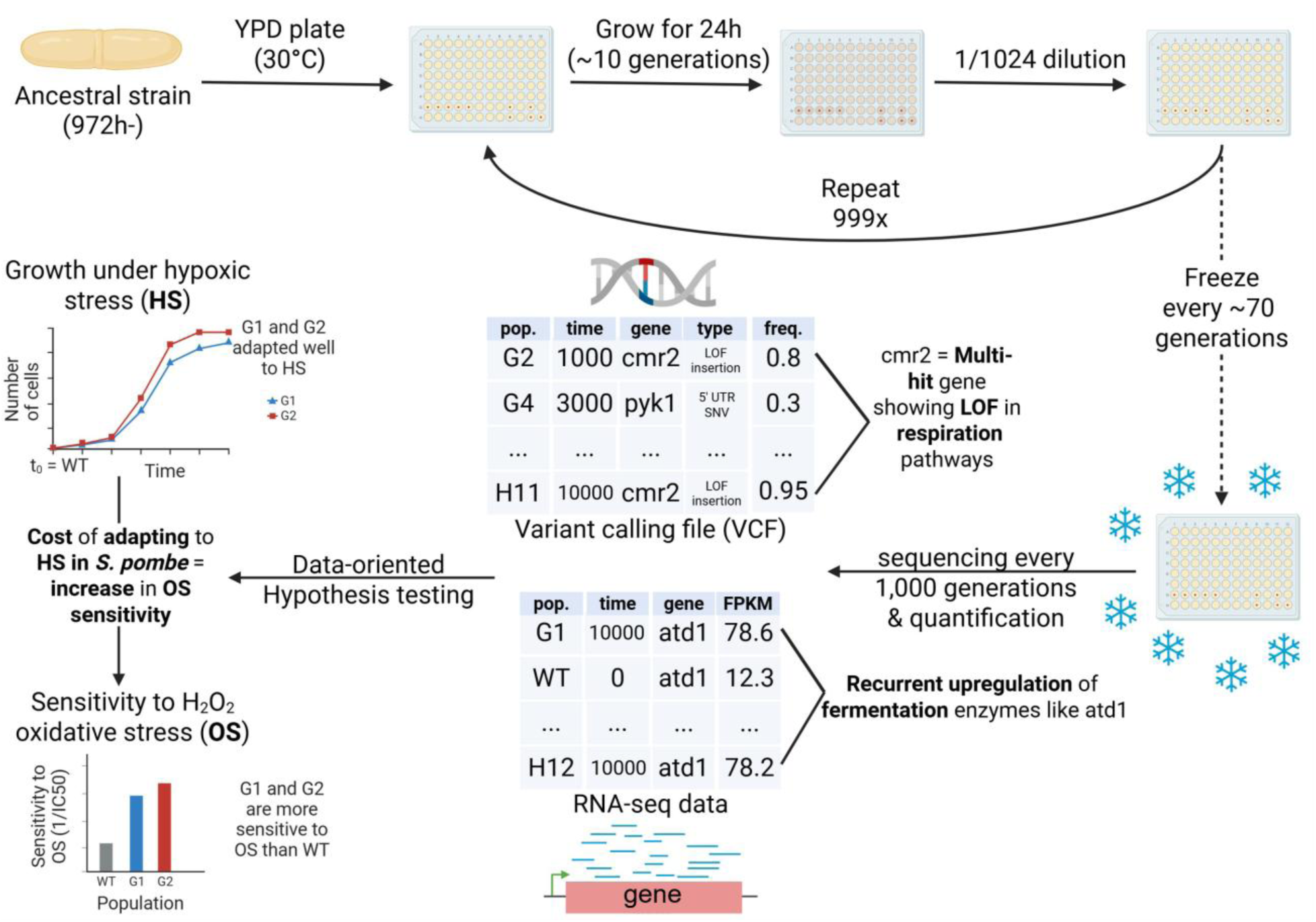
Experimental approach. We evolved 15 *S. pombe* populations at 30°C in lidded YPD 96-well plates for 10,000 generations, collecting and freezing samples approximately every 70 generations in 27% glycerol at -80 °C and sequenced them every 1,000 generations. These populations were part of a larger experimental evolution project, which explains why they occupy two rows of the 96-well plates and were examined regularly for contamination (**Methods**). Because the experimental conditions yielded a doubling rate of 10 generations per day, we performed a 1/1024 serial dilution every day (1/2^10^), which can be done by two consecutive 1:32 dilutions with 96-well plates. This yielded ∼100 time points or samples from which we sequenced DNA to identify variants. We also sequenced RNA at the initial (WT) and final time points to quantify gene expression changes. To find targets of selection at the genomic levels, we identified multi-hit genes, i.e. the genes that were recurrently hit by fixed mutations (frequency >= 75%) that are not synonymous (indels, missense, or nonsense) in multiple populations. At the transcriptomic level, we identified genes that were differentially expressed across several evolved populations compared to WT or to populations not exhibiting a change in a phenotype of interest, e.g. sensitivity to oxidative stress (OS). These data allowed us to generate hypotheses about adaptation in the 10 populations surviving, i.e. G1-G5, G9, H9 and H11-12, which we tested experimentally. For instance, in the figure, G1 (blue) and G2 (red) both adapts to hypoxic stress (HS) at the cost of becoming more sensitive to OS.

To determine whether the evolved populations adapted to these conditions, we measured the fitness at the initial and final time points. We serially propagated each population and the ancestral strain separately in a mixture with a *S. cerevisiae* reference strain expressing green fluorescent protein (GFP) for 3 days. We used a flow cytometer to measure the frequency of the dark *S. pombe* populations and the green *S. cerevisiae* strain daily, and assessed fitness based on changes in frequency (**Equation 1 in Methods**). The evolved populations are all fitter than the ancestral strain (**Supplementary Fig. 1**) as observed in other experimental evolution studies^3,6^. However, the fitness gains of the *S. pombe* lineages after 10,000 generations of evolution are significantly smaller than those of *S. cerevisiae*’s haploid populations in a similar rich-media hypoxic environment (**Supplementary Fig. 1**). This slower rate of fitness gain is not due to *S. pombe* being ‘pre-adapted’ to this environment, as the ancestor *S. cerevisiae* strain used in a previous study is much fitter than the ancestor *S. pombe* strain used here. We presume that this is the result of *S. cerevisiae* having higher glycolytic and fermentation rates and being able to more readily adapt to the experimental conditions through a more diverse set of stress response pathways as a result of having explored many anaerobic niches throughout its evolutionary history^32,34,35,45^. *S. pombe*, on the other hand, is a poor fermenter in anaerobic conditions^32^ and has a less robust hypoxic stress response^46^. Together, these factors may partly explain differences in adaptation rates under the same environmental conditions, as each species may evolve through different adaptation strategies with different associated costs.

### Differences in the genome architecture partially explain the different mutation spectra of *S. pombe* and *S. cerevisiae*

To uncover potential differences in adaptation strategies between the two species and the molecular mechanisms underlying them in the evolved populations, we performed whole-population, whole-genome sequencing every 1,000 generations on the 10 *S. pombe* populations. We successfully grew cells from 93 out of the 100 corresponding stored samples and sequenced these time points using Illumina NovaSeq paired-end sequencing (mean coverage ∼40X). The remaining 7 time points were presumably lost during storage and were not associated to any specific population or particular archival date. We then called single-nucleotide variants (SNVs) and indels using Varscan2.4.6^47^. Because contamination, sequencing errors and low-complexity regions can introduce variant calling errors, we applied some filters to the called variants ^48,49^ while leveraging the time series for call consistency (**Methods**).

To validate the accuracy of the called variants, we measured the substitution rates and fixed indel bias, e.g., the fixed insertion-to-deletion ratios, in each *S. pombe* population. We measured a fixed insertion-to-deletion ratio ranging from 2.11 to 6.33 (median = 4.43, µ= 4.32, and σ= 1.22; **Supplementary Fig. 2**), which is consistent with the insertion preference observed in previous *S. pombe* genomic analyses ^37^. As for the substitution rate, we measured an average of 1.91E-10 fixed substitutions per site per generation (median = 1.72E-10, σ= 6.76E-11), which is similar to results from mutation accumulation experiments in *S. pombe* (2.00E-10 in Farlow et al. 2015) and within the range of *S. cerevisiae*’s substitution rate estimates (1.67 ± 0.04E-10 in Zhu et al. 2014 and 3.30± 0.80E-10 in Lynch et al. 2008)^37,50,51^.

Although both species accumulate substitutions at a similar rate, their genome architectures differ. *S. cerevisiae* is a domesticated post-WGD yeast species, contrary to *S. pombe* ^52–55^. Furthermore, the extent of coding regions in *S. pombe*’s genome (∼60%) is smaller than in *S. cerevisiae* (∼70%) ^34,37,56,57^. Under a null model of evolution in which all loci have the same chance of being hit by substitutions, the differences in the mutation spectra of these species should reflect the differences in genome architecture. Deviations from this expectation would suggest the presence of selection in certain loci and a preference for specific types of mutations. Thus, we compared the frequency of different variants observed in the 10 *S. pombe* haploid populations to the one observed in *S. cerevisiae* haploid populations’ experimental evolution^3^ (**Fig. 2A**).

**Figure 2.**
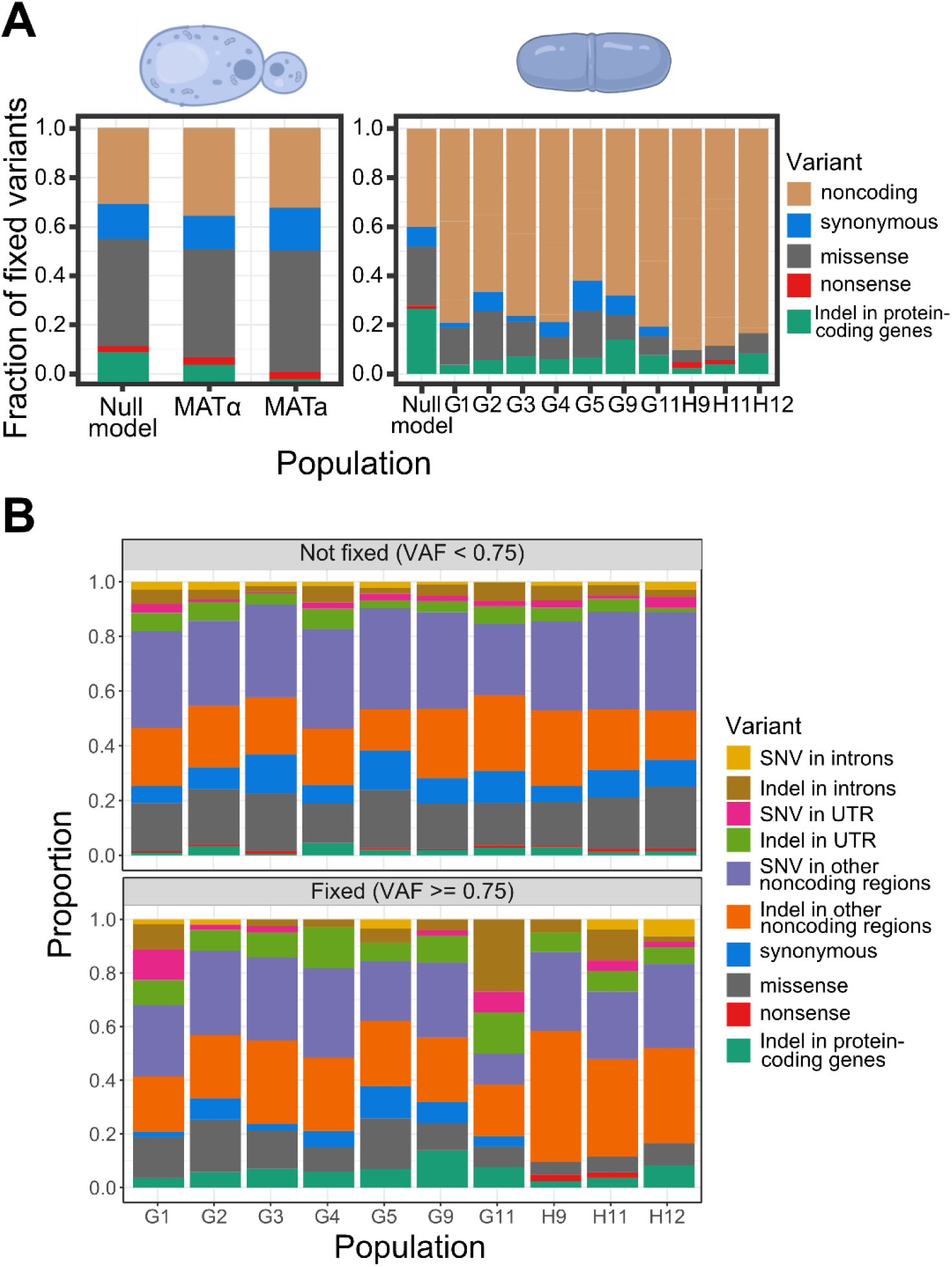
Comparison of the fixed variant spectra between the budding and the fission yeasts and the fixation bias in *S. pombe* evolved populations. A) The fixed variant (allele frequency >= 75%) spectrum in *S. cerevisiae*’s haploid populations after 10,000 generations of experimental evolution (left, data from Johnson et al. 2021) compared to *S. pombe*’s (right). The experimental evolution protocol is the same for both species and is described in Johnson et al. 2021^3^. The null model is obtained by calculating the expected number of fixed variants in each category based on the species’ evolution rate, genome size, genome architecture, and the proportion of single-nucleotide mutations derived from the genetic code and associated with a specific category of protein-coding region mutations (**Equations 2 and 3 in Methods**). B) The upper part of this panel represents non-fixed variants, while the bottom part represents the fixed variants in each of the 10 evolved *S. pombe* populations.

The fixation of variants in the noncoding regions of *S. pombe* is higher than expected by the null model, suggesting the predominance of positive selection in noncoding regions or purifying selection in coding regions. The fission yeast populations mainly evolved through mutations in non-coding regions with a bias toward fixing indels and not fixing coding regions SNVs (**Fig. 2B**), which is different than what has been reported for the budding yeast^3^. To test if the stronger fixation bias of indels in non-coding regions compared to protein-coding genes reflects a signal of purifying selection in protein-coding genes, we compared the indel rate, i.e., the number of indels per kb, based on whether the indel locus is coding or non-coding and whether the indel changes the DNA reading frame or not (**Supplementary Fig. 3**). In protein-coding genes, controlling for increased likelihood of shorter indels, we find that indels that do not cause frameshifts are enriched compared to the ones that do (**Supplementary Fig. 3**). In contrast, indels in non-coding regions do not exhibit such selective constraints on their length. This is similar to the observed frequencies of polymorphisms in humans and budding yeast^58^ and suggests stronger purifying selection in protein-coding genes. This and the paucity of fixed coding mutations compared to *S. cerevisiae* populations suggest that the genetic context of *S. pombe* is less amenable to tinkering through loss-of-function mutations. One hypothesis for this may be that the whole-genome duplication event in *S. cerevisiae* provided more genetic redundancy, which has been suggested to increase evolvability through coding mutations or through rapid changes in functional gene content^30,59^ (see Discussion). An additional explanation is that *S. pombe* has a much higher essential gene load than S. cerevisiae (26.1% vs 17.8%)^60^. Altogether, these results show that the differences in the mutation spectra of *S. cerevisiae* and *S. pombe* are only partially explained by their genome architectures, revealing signatures of selection in both coding and non-coding regions.

### The genomic targets of adaptation in *S. pombe* are genes involved in the response to hypoxic and oxidative stresses

Although the *S. pombe* mutation spectrum revealed a signal of purifying selection in protein-coding genes, we cannot exclude the presence of loci evolving under positive selection. However, we may expect the targets of adaptation to vary between fission and budding yeasts due to differences in their evolutionary histories and genetic architectures. To detect such loci in 205 *S. cerevisiae* populations, Johnson et al. (2021) looked for signals of convergent evolution. More precisely, they defined a “hit” as a fixed variant that is not synonymous (missense, nonsense and indels), and a multi-hit gene as a gene having hits in multiple populations. This enabled the identification of 1,092 genes with hits in at least 3 out of 205 populations. Among these genes, they identified a set of 189 genes with hits in at least 6 populations that were enriched in functions related to the adenine biosynthesis pathway, the mating pathway, and negative regulators of the Ras pathway. These signals of parallelism at the levels of biosynthetic and signaling pathways were driven by LOF mutations, which were primarily frameshift deletions and nonsense mutations.

Using the same approach with a minimum threshold of 2 out of 10 populations in which a gene has a hit, we identified 19 multi-hit genes with hits in either the cis-regulatory region or the coding region (**Table 1**). We note, however, that 10 indels were found repeatedly in many populations (but not all). These may indicate remarkable genetic-level nucleotide convergence (or rather strong mutational biases), or they may signal the presence of sequencing artifacts or contamination. Upon inspection of these indels, we found that they were in homopolymeric tracts and overwhelmingly had consistent allele frequency trajectories that, in isolation, would have indicated a *bona fide* mutation (see **Methods** for details in how we characterize these). We thus include them in our analyses but refrain from deriving any biological conclusions from these. If they represent true mutational biases or signals of selection, we believe they warrant further study.

**Table 1.** Multi-hit genes and their LOF mutations. The gene annotations are based on Pombase data^40^.

| Gene symbol/<br>systematic ID<br>(Number of<br>populations with<br>a hit (PH)) | Function(s)<br>(biological<br>process) | Gene category<br>(Link with the<br>response to HS<br>and OS) | Characterization of the hit(s)<br>(Gene product truncation or<br>potential LOF) |
| --- | --- | --- | --- |
| *bmc1*/<br>SPBC2A9.10<br>(2) | Bin3 family RNA<br>methyltransferase<br>(snRNA<br>metabolic<br>process) | Others | 1 frameshift insertion in both G1 and G3 (Truncation of 14 out of 268 residues) |
| *bop1*/<br>SPAP32A8.03c<br>(2) | ubiquitin-protein<br>ligase E3<br>(protein<br>degradation) | HS-related<br>(Ubiquitin-dependent<br>protein<br>degradation is<br>ATP-dependent<sup>61,62</sup>.<br>The response to<br>HS requires<br>specific E3<br>ligases that<br>activate sterol<br>regulatory<br>elements (SRE)<br>and the decrease<br>of the activity of<br>other E3 ligases<br>that do not<br>contribute to the<br>HS response,<br>e.g. Ubr1<sup>63–65</sup>) | 1 frameshift insertion in both G11 and H12 (Truncation of 475 out of 513 residues) |
| ***clg1/***<br>**SPBC1D7.03**<br>**(2)** | Cyclin-like protein involved in autophagy (signaling, molecular activity regulation and autophagy) | HS- and OS-related (in *S. pombe*, *clg1* regulates autophagy<sup>66</sup>. Autophagy plays an important role in the response to HS and OS by recycling biomolecules and removing damaged cell components<sup>67,68</sup>) | 1 indel in the 5'UTR of the populations G2 and G3 |
| ***cmr2/***<br>**SPAC56F8.02**<br>**(2)** | acetyl-CoA biosynthesis (respiration) | HS- and OS-related (Acetyl-CoA is involved in the first step of the TCA cycle and respiration enzymes are involved in energy production and ROS removal<sup>69</sup>) | 1 missense mutation in the AMP-dependent synthetase/ligase domain<sup>40</sup> in G1<br>1 nonsense SNP in H11 (Truncation of 174 amino acid residues in the Dip2-like domain<sup>40</sup> out of 1517 protein residues) |
| ***ctf18/***<br>**SPBC902.02c**<br>**(4)** | Ctf18 RFC-like complex subunit Ctf18 (DNA replication) | HS-related (Upregulated under HS<sup>67</sup>) | 1 indel in the first intron of the gene in the populations G3, G5, H9, and H11 |
| ***dep1/***<br>**SPBC21C3.02c**<br>**(2)** | Rpd3L histone deacetylase complex subunit (chromatin organization and transcription regulation) | HS-related (Upregulated under HS<sup>67</sup>. *dep1Δ* strain is more sensitive to hydrogen peroxide<sup>70</sup> than WT) | 1 indel in 3'UTR in the populations G2 and G11 |
| ***fet4/***<br>**SPBP26C9.03c**<br>**(2)** | plasma membrane iron/zinc ion transmembrane transporter | HS- and OS-related (Iron cofactors support major biochemical processes like | 1 indel in 5'UTR in the populations G2 and H11 |
|  | (inorganic ion homeostasis and transmembrane transport) | the heme biosynthesis, TCA cycle, electron transport chain, translation, and DNA repair<sup>71)</sup> |  |
| ***fhn1/***<br>**SPBC1685.13**<br>**(3)** | Eisosome assembly protein (membrane organization) | HS-related (Upregulated during the response to hydrogen peroxide<sup>72)</sup>) | 1 indel in 3'UTR in the populations G3 and H11.<br>1 indel in 3'UTR in the populations G9 |
| ***iss9/***<br>**SPBC2A9.11c**<br>**(2)** | SAC3/GANP/TH P3 family protein (transcription or RNA surveillance) | Others | 1 frameshift insertion in the populations G1 and G5 (Truncation of 315 out of 395 residues) |
| ***pfl2***<br>**SPAPB15E9.01c**<br>**(3)** | cell surface glycoprotein, flocculin Pfl2 (cell adhesion) | HS- and OS-related (Flocculation and adhesion enhance survival by protecting against stressful conditions<sup>73,74)</sup>) | 5 missense hits in G3: chrI_A3989282G (R->G), chrI_G3989285A (A->T), chrI_G3989294A (G->S), chrI_G3989384A (G->S), chrI_G3989390A (A->T)<br>4 missense hits in G5: chrI_G3989877C (R->T), chrI_G3989879A (G->R), chrI_A3989878T (R->S), chrI_G3989877C (R->T)<br>*Pfl2 is a disordered protein so it has many S, R and T residues<sup>40,75)</sup><br>2 indels in 5'UTR in the populations G1 and G5 |
| ***pfl5/***<br>**SPBC1289.15**<br>**(3)** | cell surface glycoprotein, flocculin Pfl5 (cell adhesion) | HS- and OS-related (Flocculation and adhesion enhance survival by protecting against stressful conditions<sup>73,74)</sup>) | 2 Y→F missense hits in G2 and H12 around the middle of the protein. 1 K→E missense hit in G5<br>The same frameshift insertion in the populations G2 and H12 (Truncation of 518 out of 1283 residues around the middle of the protein).<br>1 deletion in G2 (Truncation of 528 out of |
|  |  |  | 1283 residues around the middle of the protein) |
| ***pof15/***<br>**SPAPB1A10.14**<br>**(2)** | Ubiquitin complex adaptor; F-box protein (protein degradation) | HS-related (Downregulated under HS<sup>67,76</sup>. Ubiquitin-dependent protein degradation is ATP-dependent<sup>61,62</sup>) | • 1 N→D missense hit in G5<br>• 1 frameshift deletion in the population H12 (Truncation of 135 out of 243 residues) |
| ***pqr1/***<br>**SPAC6B12.07c**<br>**(3)** | SPX-RING-type ubiquitin-protein ligase regulating phosphate transport and homeostasis (protein degradation and transmembrane transport) | HS-related (Upregulated during the response to hydrogen peroxide<sup>72</sup>. Excess of intracellular phosphate and polyphosphate is associated with improper autophagy-dependent proteolysis in vacuoles<sup>77</sup>) | • 1 SNV in 3'UTR in G9, G11 and H11 |
| ***prr1/***<br>**SPAC8C9.14**<br>**(2)** | stress-responsive DNA-binding transcription factor Prr1 (transcription regulation and response to oxidative stress) | OS-related (*prr1* is a positive regulator of the OS response. It binds *pap1* as part of a positive regulation of transcription, and it regulates *ctl1*, *srx1*, *trr1* during the cellular response to OS<sup>78,79</sup>) | • 1 P→Q missense hit in the signal transduction response regulator/receiver domain in H12<br>• 1 frameshift insertion in the population H11 (Truncation of 517 out of 540 residues) |
| ***pyk1/***<br>**SPAC4H3.10c**<br>**(5)** | pyruvate kinase (carbohydrate metabolic process) | HS- and OS-related (The gene product catalyzes the last step of glycolysis. This | • 1 missense hit V→L in the barrel domain in H11<br>• 2 SNVs in 3' UTR in G1<br>• 1 SNV in 3' UTR in H12<br>• 1 deletion in the promoter (5' UTR) in G3 |
|  |  | influences the fermentation-respiration balance and thus biomass production and stress resistance<sup>35)</sup>) | 1 deletion in 5' UTR in G4 |
| ***pzh1*/**<br>**SPAC57A7.08**<br>(2) | serine/threonine protein phosphatase (signaling) | HS- and OS-related (*pzh1* is a positive regulator of the response to OS<sup>80</sup>. It is also upregulated under HS<sup>67)</sup>) | 1 deletion of 2 residues (positions 73 and 74) in the N-terminal disordered region<sup>40</sup> in both G5 and G9 |
| ***rpp1*/**<br>**SPAC3A12.04c**<br>(2) | RNase P and RNase MRP subunit p30 (ribosome biogenesis and tRNA metabolic process) | Others | 1 SNV in the fourth intron of the gene in populations H11 and H12 |
| ***seb1*/**<br>**SPAC222.09**<br>(2) | poly(A)site selection protein (mRNA metabolic process and snoRNA metabolic process) | Others | 1 missense T→I hit in the RNA recognition motif domain in G1<br>1 missense K→N hit in the CTD-interacting domain in G2 |
| ***ubp9*/**<br>**SPBC1703.12**<br>(2) | ubiquitin C-terminal hydrolase Ubp9 (protein degradation and vesicle-mediated transport) | HS- and OS-related (*ubp9Δ* strain is more sensitive to hydrogen peroxide<sup>70</sup>. Protein degradation and vesicle-mediated transport enable the recycling of biomolecules under stress conditions<sup>61,62)</sup>) | 1 frameshift insertion in the populations G3 and G9 (Truncation of 265 out of 585 residues, including most of the ubiquitin carboxyl-terminal hydrolase domain) |

The presence of hits in multi-hit genes was more recurrent than expected under a null model where variants are fixed at the same rate across all the genes (**Methods**; Two-sided Wilcoxon rank sum p-value < 0.05; **Supplementary Fig. 4**). This signal of positive selection is also consistent with the fixation of these hits (**Fig. 3**) and the fixation of the same hit at different times in multiple populations (**Table 1**; **Fig. 3**), though we note some of multi-hit genes may be due to mutational biases.

**Figure 3.**
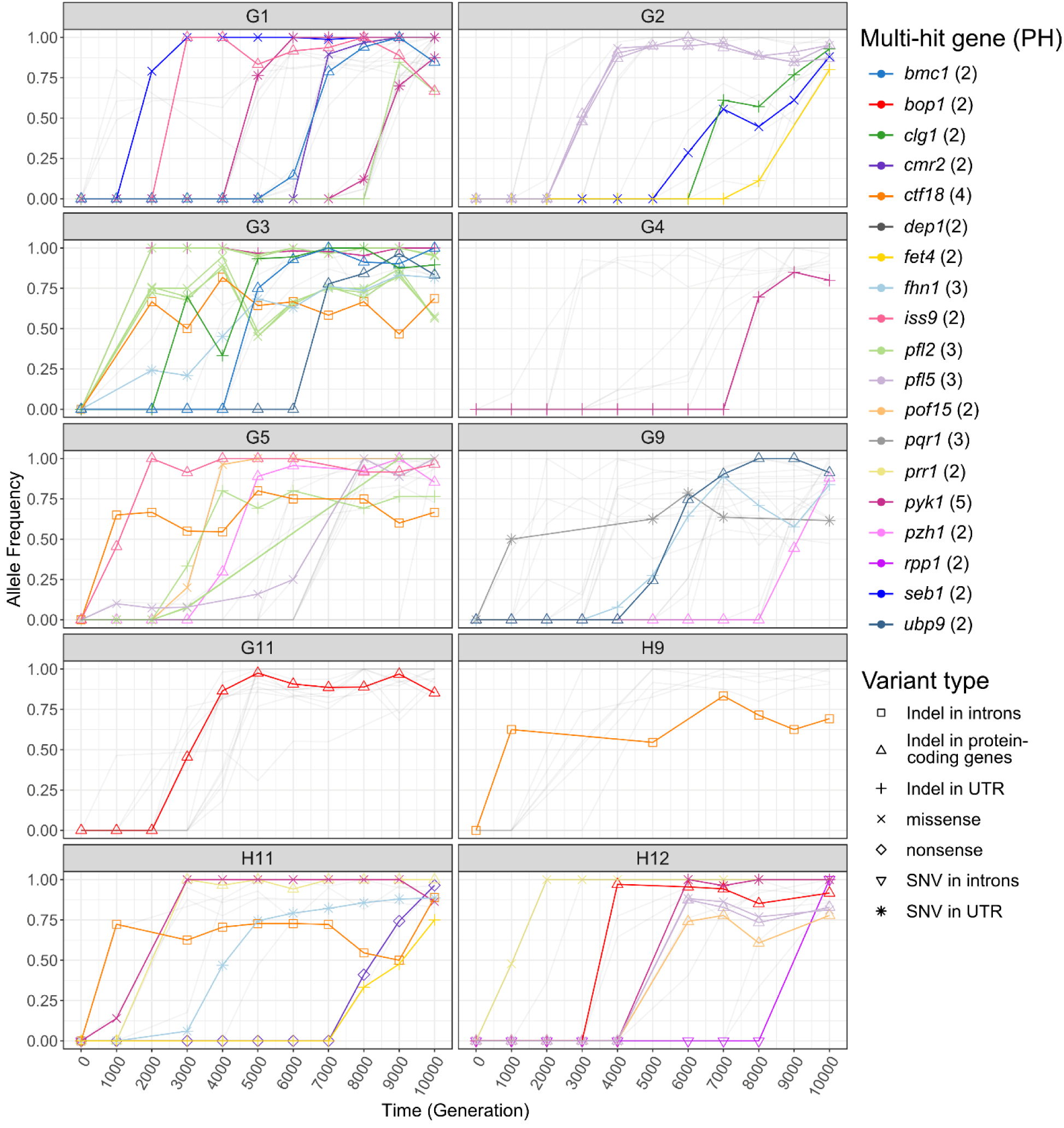
Time series of the hits’ allele frequencies in evolved *S. pombe* populations and multi-hit genes. Time series of the hits’ allele frequencies in the *S. pombe* populations. We defined a “hit” as a fixed genic variant that is not synonymous (missense, nonsense and indels) and a multi-hit gene as a gene having hits in multiple populations. In each population, variants that were never fixed or that are not located in multi-hit genes are represented by more transparent lines. In the legend, PH represents the number of populations in which the gene is hit.

We also found evidence for adaptation through LOF mutations in the 10 evolved *S. pombe* populations, although these mutations are primarily frameshift insertions rather than deletions. This is consistent with a different indel bias than in *S. cerevisiae*^3^ (**Table 1**; **Fig. 2**) and a preference for insertion, which we previously highlighted (**Supplementary Fig. 2**).

Most of the *S. pombe* multi-hit genes are involved in the response to hypoxic stress (HS) and sometimes to oxidative stress (OS) (**Table 1**), which may be due to growth in an unshaken and lidded environment. Adapting to HS requires compensating for the energy deficit caused by respiration deficiency through fermentation and removing the associated reactive oxygen species (ROS)^32^. Further, the increase in ethanol or acetate production through fermentation can hinder mitochondrial membrane integrity and functions, leading to ROS production^45,81–85^. This is consistent with fixed mutations in genes involved in the removal of damaged cell components and ROS like *clg1*, *pqr1*, *pzh1* and *ubp9*. Fermentation is also known to yield smaller energy and biomass outputs than respiration ^32,34^, which can affect the synthesis of redox metabolism components. Finally, *S. pombe* is known to be sensitive to polypeptones included in the growth media of the experiment (due to cell lysis), and it is thought that the respiratory Coenzyme Q (CoQ10) plays a role in promoting this trait^86,87^. Under the same conditions, *S. cerevisiae*, being petite-positive, can rely on fermentation along with a robust antioxidant response to mitigate the ROS produced by fermentation and hypoxic stress^45^, which may explain why such mutations are not typically observed in budding yeast experimental evolution studies.

Another piece of evidence supporting the importance of energy- and ROS-related processes in this environment is the fact that the strongest signal of convergent evolution is in the lower glycolysis gene *pyk1*, which had fixed mutations in 5 populations. This gene encodes for the enzyme involved in the synthesis of pyruvate, which influences respiration and fermentation as it can be transformed into Acetyl-CoA or into acetate and ethanol, the fermentation products^35^. In the population H11, the glycolytic flux going towards respiration seems to be compromised by a fixed LOF mutation in *cmr2* (**Fig. 3**), which encodes for the enzyme converting pyruvate into Acetyl-CoA. The fixation of this LOF mutation might be related to a benefit in re-orienting the glycolytic flux mostly towards fermentation. In the same population, we also observed the fixation of a LOF mutation in *prr1*, which encodes a transcription factor involved in the response to OS. Because *prr1* is not known to be directly activated by HS, there might be other OS response genes involved in ROS removal under HS, and *prr1* might not be needed unless ROS levels exceed the removal capabilities of these genes. For instance, in *S. pombe,* the transcription factor Sre1 is activated under low oxygen levels and activates genes involved in ROS removal or protein catabolism like *ccs1* and *SPAC23C11.06c*^67,88^. The benefit of inactivating *prr1* while respiration is limited under HS might thus be related to conserving energy and cell resources. However, losing such response regulators and relying on fermentation may make *S. pombe* more sensitive to OS.

### Most evolved *S. pombe* populations are more sensitive to oxidative stress

As we observed fixed LOF mutations in genes modulating OS response and the glycolytic flux towards respiration, we hypothesized that evolved populations might be more sensitive to OS. Populations adapting to hypoxic stress or to polypeptone-mediated lysis may exhibit decreased respiration rates, resulting in reduced energy and biomass production^32^. This could affect the synthesis of the components of the redox metabolism, e.g. glutathione (GSH) and ubiquinone (CoQ10)^69,89^. These enzymes and the ones involved in the respiratory chain rely on the biomass produced by glycolysis and the TCA cycle as well as ions like iron and sulfur for their functions^32,89–91^. Consistent with this, we observed the presence of hits in the iron transporter *fet4* in multiple populations, which may compensate for respiration deficiency. Indeed, iron can help capture or transport the low amount of oxygen available, and iron is a critical component of the respiratory chain enzymes. Moreover, one *S. pombe* evolved population harbors a LOF insertion in *prr1* (**Table1**, **Fig. 3**), an important transcription factor involved in the response to oxidative stress^79^. As *prr1* activates the catalase *ctt1*, the sulfiredoxin *srx1*, and the thioredoxin *trr1*^78^, a fixed LOF mutation in this gene suggests that evolving under HS can decrease *S. pombe*’s ability to manage OS, which is consistent with observations in the cousin species *S. japonicus*^32^. These species likely rely on fermentation to adapt to hypoxic conditions, which may expose them to more ROS^45^. Altogether, these observations led us to believe that the evolved *S. pombe* populations are more sensitive to OS than WT as a result of their adaptation to HS.

To test this, we used hydrogen peroxide as a source of oxidative stress during a growth assay (**Methods**) and compared the OS sensitivity of the WT to the OS sensitivity of the 10 evolved populations (**Fig. 4A**).

**Figure 4.**
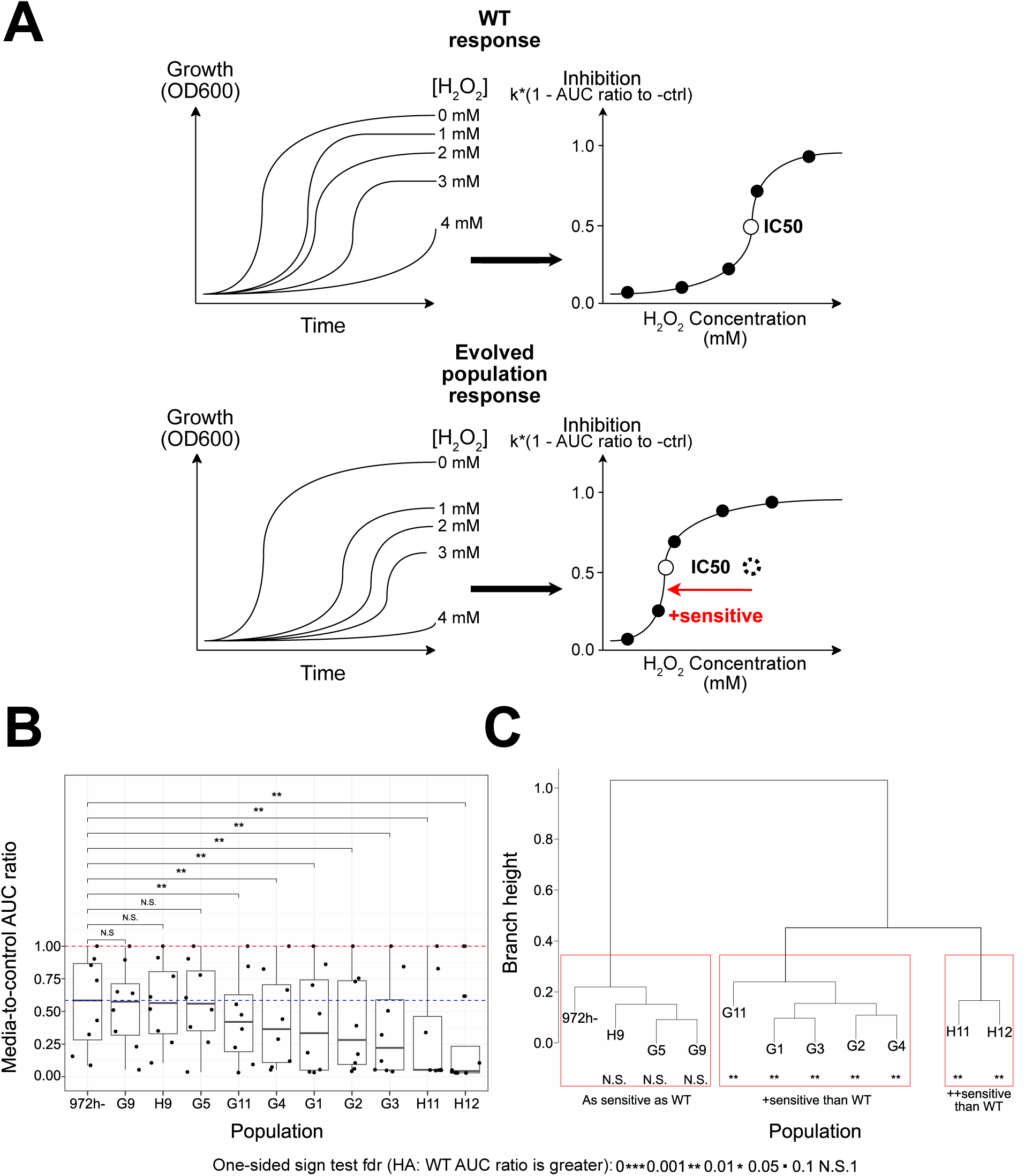
Comparison of the sensitivity levels to H_2_O_2_ between *S. pombe* WT and evolved populations. A) Schematic illustrating the expected difference in sensitivity between WT and evolved populations. We inferred the degree of inhibition at each concentration of H_2_O_2_ based on the expression in Equation 4 and the sensitivity of each population based on Dose-Response analysis (**Methods**). The IC50 recapitulates the level of sensitivity of a population to hydrogen peroxide. Lower IC50 values can be interpreted as higher sensitivity to H_2_O_2_, which is what we expected for the evolved populations. B) The ratio between the AUC of a population at a certain H_2_O_2_ concentration + YPD and the AUC in YPD without H_2_O_2_. Each point is a different concentration. This is another metric of sensitivity with an expected maximum of 1 (red dotted line). The blue dotted line represents the median sensitivity of the WT. The WT AUC ratio is compared to each evolved population using a sign test on the ranks. The one-sided adjusted p-values (false discovery rates; alternative hypothesis=“WT AUC ratio is greater than evolved populations’”) of this test are encoded as follows: *** from 0 to 0.001 exclusively, ** from 0.001 to 0.01 exclusively, * from 0.01 to 0.05 exclusively, “.” from 0.05 to 0.1 exclusively and N.S. otherwise, which is the acronym of “non-significant”. C) Red boxes represent the population clusters, which are based on the AUC ratios.

The evolved populations clustered into 3 groups: 3 out of 10 populations had a similar sensitivity to H_2_O_2_ as the WT, 5 out of 10 populations were more sensitive than WT, and 2 out of 10 evolved populations were much more sensitive than the WT strain, including H11, the population that lost the OS response regulator *prr1* (**Fig. 4B** and **4C**). This increased sensitivity to OS after adapting to HS has not been observed in *S. cerevisiae*, which may be explained by contingency, i.e. more exposure to anaerobic environments during its evolutionary history and a better redox metabolism^32,34^. However, it is reproducible in most of the evolved *S. pombe* populations, and has likewise been observed in related species, including wild *S. japonicus*^32^ (see Discussion).

### Transcriptomics analysis reveals many genes involved in the response to HS and downstream effects explaining the sensitivity to OS in multiple populations

The sensitivity to OS developed after adapting to HS was revealed by convergent evolution at the genomic level, i.e. multi-hit genes. However, identifying transcriptomic differences between more sensitive and WT-like populations could help reveal genes driving this evolutionary outcome and explain how point mutations and indels contribute to the evolution of these traits through transcriptomic regulation. Moreover, most of the mutations in the pyruvate kinase (*pyk1*) locus are located in the untranslated regions flanking the gene, which contains cis-regulatory elements (CRE) like the promoter (**Supplementary Fig. 5**). Hits in these loci could explain the adaptation to HS through changes in transcript levels through cis-regulatory and trans-regulatory effects^92–96^. These questions thus led us to investigate the role of transcriptomic changes in the responses to HS and OS. To do so, we performed RNA-seq of the WT ancestor and the final evolved populations, and compared the transcriptomic profiles of populations more sensitive to H_2_O_2_ than WT with those of populations as sensitive as WT (**Methods**).

Consistent with the need to compensate for biomass and energy deficit under HS due to limited respiration^32^, most evolved *S. pombe* populations upregulated genes involved in heme metabolism/transport, the transport of biomolecules like amino acids or hexose, lipid metabolism, fermentation, and ion transport (**Fig. 5, Supplementary Table 1**).

**Figure 5.**
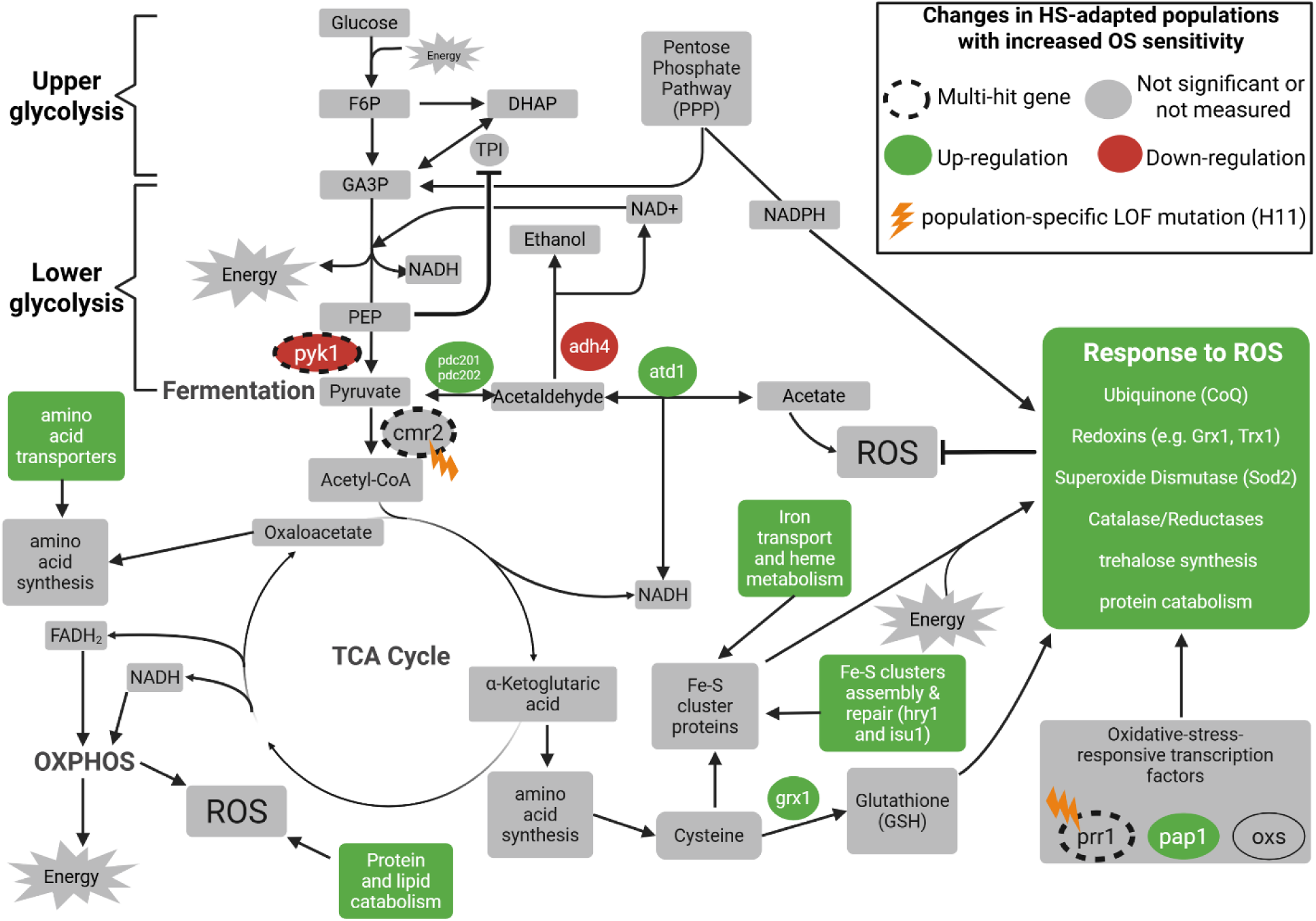
Fission yeast cellular responses to hypoxic stress in populations with increased OS sensitivity. Transcriptomic changes are determined by comparing the populations with increased OS sensitivity to populations with similar OS sensitivity than WT while genomic changes are obtained by comparing these evolved populations to WT (**Methods**). The green fill represents an upregulation while the red fill represents a downregulation. When processes are color filled, they represent cases where some genes involved in those processes were upregulated. The grey fill represents molecules or processes that were not associated with a gene with significant effect or not measured (no data). The orange lightning symbol represents LOF hits in H11, the only population with such variants in several multi-hit genes (dotted line stroke).

Another way the fission yeast populations could respond to HS is by improving the electron transport chain (ETC) to leverage the limited amount of oxygen available. The upregulation of the heme transporters *str3* and *shu1* in many evolved populations supports such a strategy (**Supplementary Table 1**). Heme can help capture and transport the small amount of oxygen available under HS and it is an essential component of the electron transport chain (ETC)^90^.

Despite the presence of transcriptomic changes explaining HS adaptation and the OS sensitivity cost, most of the differentially expressed genes were different than the targets of selection identified at the genomic level, except for *pyk1*, *pzh1* and *ubp9*. For instance, the multi-hit gene *clg1*, which is a positive regulator of autophagy^40,66^, is not differentially expressed between populations with different OS sensitivity, but 22 other genes involved in autophagy exhibit an expression level increase from 54.4% to 229% (**Supplementary Table 1**) and they are not mutated. Autophagy might be one of the few pathways that contribute to cross-resistance to both HS and OS by recycling biomolecules and removing damaged cell components^67,68^. This is consistent with the upregulation of the multi-hit gene *pqr1*, the ubiquitin-protein ligase regulating phosphate transport and homeostasis, such that autophagy is not compromised in vacuoles^77^. Another process linked to cross-resistance to multiple stresses is vesicular transport^97,98^. One of its genes, i.e. *ubp9,* is one of the only three multi-hit genes to be differentially expressed in evolved *S. pombe* populations that are more sensitive to OS than WT (log2FC = 0.867; FDR<0.05), suggesting a moderate but significant link between vesicular transport and adaptation (**Supplementary Table 1)**. As for the multi-hit gene *pyk1*, it is downregulated by 36.5% in populations with increased OS sensitivity (FDR<0.05; **Supplementary Fig. 6**). Adaptation to HS may require decreasing the glycolytic flux allocated to respiration^32^ and change the respiration/fermentation balance. This is consistent with the decrease of Acetyl-CoA synthesis through the downregulation of the Acetyl-CoA ligase *pcs60* (53.7% decrease; FDR<0.05) and selected LOF mutations in *cmr2*, which may trigger the ETC optimization responses previously described as a compensatory mechanism. Similar to mutations observed in *S. japonicus*, the LOF mutations in lower glycolysis and Acetyl-CoA synthesis might also contribute to OS sensitivity in *S. pombe* as it would decrease the biomass and energy output of respiration, which is used for the synthesis of redox metabolism proteins^69,89^. The last differentially expressed multi-hit gene is *pzh1* (log2FC = 0.752; FDR<0.05), which regulates the activity of the plasma membrane H+-ATPase^99^. This is consistent with the need to increase energy production to compensate for respiration deficiency in hypoxic conditions.

Apart from these three multi-hit genes, transcriptomic changes shared by populations with increased OS sensitivity involved genes that were not mutated at the genomic level (**Supplementary Table 1; Table 1**). This is the case for many upregulated genes involved in fermentation, e.g. *pdc201/202* and *atd1*, and in the synthesis of ETC components and their iron-sulfur clusters, e.g. *cbp3-4, cbp6, coq9, sdh3, sdh8*, *qcr7-10*, *fxn1* and *isu1*. The lack of overlap between selection targets at the genomic and transcriptomic levels suggests that the overall changes in expression levels shared by many evolved populations are driven by trans-regulatory rather than by cis-regulatory effects. However, interpreting metabolic fluxes from transcriptomics data remains challenging and further exploration into the mechanisms affected by this rewiring is needed to reveal more deeply how they relate to adaptation in our environment.

## DISCUSSION

Over short periods of time, experimental evolution is a powerful tool to decipher molecular pathways under selective constraints^3,11^. An important result from studies exploiting asexual laboratory evolution is that evolution is usually not limited by mutation: adaptation usually proceeds rapidly and is often dominated by large-effect mutations. In many cases, gene content variation can occur (via ploidy changes), and mutations frequently inactivate gene function. Perhaps the most interesting result from many of these studies is that laboratory evolution can frequently recapitulate evolutionary outcomes from wild, uncontrolled evolution. Whole-genome duplication, loss-of-function mutations, the rise of mutators, stable co-existence, and like here the change of carbon flow towards increased fermentation, are all phenomena thought to have occurred in various species at various times during their evolutionary history, but also notably in cancer cells^11,100–102^. The fact that cancer cells are also evolving populations suggests that experimental evolution in different single-cell organisms can reveal fundamental principles of the molecular basis of adaptation. For instance, the pervasiveness of parallel evolution in the tree of life could be explained by deep conservation of interconnected biological pathways.

Our study extends the discovery of shared molecular mechanisms of adaptation across scales in diverse biological systems. For instance, *S. pombe* shares a distinct preference for insertions as a mutational bias with *C. elegans*^38^,. On specific selection targets, mutations that lead to upregulation of iron metabolism, autophagy regulators and vesicle transporters, and that lead to rewiring of the glycolytic flux are common observations in populations adapting under iron and oxygen limitation^104,105^ (such as tumors). The fact that the orthologs *pykF* and *pyk1* in *E. coli* and *S. pombe*, respectively, are both recurrently hit across multiple populations illustrates the conservation of some selection targets at the gene level, even across distant species. Finally, parallel evolution within model systems is frequent and has been observed in nearly all experimental evolution studies to date, but the precise observed trait under selection varies (examples in ^6,106^ and reviewed in^107^).

Despite all these common evolutionary mechanisms and outcomes, disparities can arise due to differences in genome architecture and contingency. One of the major findings in evolutionary dynamics is that low-fitness populations typically adapt faster than high-fitness populations. This observation has been seen in several species, and in several environmental contexts^3,6,11^. In the evolution environment of our study, the ‘ancestor’ *S. cerevisiae* is much fitter than its *S. pombe* counterpart. Yet, the fitness gains by our *S. pombe* populations are relatively modest (**Supplementary Fig. 1**). We propose that one explanation for this ‘evolvability’ is the effect that we set out to study in the first place, which is evolutionary contingency. Indeed, in a previous study, it was found that the identity of specific molecular pathways of gene deletions had a strong impact on fitness gain after evolution^13^.

Further supporting this, the most frequent targets of adaptation in the past *S. cerevisiae* study were genes involved in adenine biosynthesis and the mating pathway, for which genomic hits have high fitness effects^3^. However, these pathways are not targets of selection in these *S. pombe* populations. Based on our understanding of the adenine biosynthetic pathways in both species, mutations in adenine biosynthesis genes were not expected in our *S. pombe* populations (which are ade+). In contrast, the *S. cerevisiae* populations were *ade2*-, which led them to accumulate a toxic intermediate that can be suppressed by the observed adenine pathway mutations after evolution. It is well known that similar effects can be observed in *ade6-*strains of *S. pombe*^108^. Similarly, mutations in the mating pathway might not be observed in *S. pombe* due to its preferential haploid lifestyle and its mating pathway is not active in rich media^23,24^.

A third pathway frequently mutated in the budding yeast laboratory evolution experiment is the Ras-cAMP pathway^3,8,109^. Observed loss-of-function mutations in *IRA1* or *IRA2* (but usually not both), relieve inhibition of Ras and lead to PKA activation ^3,110^. Activation of PKA ultimately leads to increased glycolysis through phosphorylation of Pyk1^111^. Our *S. pombe* populations never mutated the *IRA1/2* ortholog (*gap1*), though we note that it is a single-copy gene in the fission yeast. More surprisingly though, are the mutations observed in *pyk1*. These mutations are not the usual loss-of-function nonsense or frameshift mutations. Instead, most mutations were found in the regulatory region of the gene, which we initially believed to be an analogous way for *S. pombe* to upregulate Pyk1 activity. However, whole-transcriptome analyses suggest that these mutations downregulate Pyk1 activity (**Fig. 5**). Interestingly, similar mutations leading to lower Pyk1 activity have been observed in wild *S. pombe* populations and in the common laboratory strain^35^, suggesting that decreasing Pyk1 activity may be adaptive. In Kamrad et al. (2020), the authors showed that a decrease in Pyk1 activity does not increase growth rate nor biomass, seemingly contradicting the strong signature of adaptation from parallel evolution. However, the authors noted that lower Pyk1 activity might be an adaptive trade-off as lower glycolytic flux increases oxidative stress resistance through higher pentose-phosphate pathway flux and NADPH production. Despite transcriptomic analyses showing a slight decrease in *pyk1* expression levels in most evolved *S. pombe* populations, they were often more sensitive to OS than WT. This is potentially explained by the fact that during our experiment, the slightly reduced glycolytic flux is mostly redirected toward fermentation, which is upregulated to compensate for the fission yeast respiration deficiency in hypoxic conditions.

Adaptation to our growth environment has led to an increased OS sensitivity for most evolved populations via multiple molecular mechanisms. First, Pyk1 activity can regulate carbon flow and, like the other mutations found in the wild, can favor fermentation rather than respiration, as our lidded and unshaking environment is likely to lead to hypoxic stress. Supporting this, we observed adaptations to HS, including LOF in enzymes responsible for the transition between glycolysis and oxidative phosphorylation, and the upregulation of the fermentation rate. This leads to an increase in intracellular ROS through higher ethanol production and a deficiency in the production of biomass and energy, which sustains the redox metabolism^34,45^. In our dataset, 3 populations in which *pyk1* is downregulated have cis-regulatory mutations around the gene, while G2 is the only population in which *pyk1* is downregulated without such mutations (**Supplementary Figs. 5 and 6**). This suggests that *pyk1* downregulation can evolve more easily through cis-regulatory effects, but it can also evolve through trans-regulatory effects. The effect of reducing Pyk1 activity here is also consistent with the other mutations we observed that appear to modulate central carbon metabolism (**Figs. 3 and 5**). Second, *S. pombe* cells are sensitive to polypeptones, a major component of our environment^86^. Mutations that lower respiration and flux through the mitochondrial electron transport chain have been shown to increase resistance to polypeptones because this sensitivity requires CoQ10^86^. Strikingly, there is also evidence in *S. japonicus* that lower glycolysis is frequently being rewired in natural populations^32^. This suggests that adaptation to HS or polypeptones in petite-negative yeast species with relatively modest fermentation rates might often come at the expense of costly carbon flow rewiring, hindering the response to OS.

Multiple genomic variants and transcriptomic changes were involved in the apparent trade-off between HS and OS, e.g. the LOF or downregulation of some genes in lower glycolysis or Acetyl-CoA synthesis, the upregulation of genes in fermentation, and the upregulation of genes in the transport and metabolism of biomolecules and ions. Among these adaptations to HS, fermentation upregulation and LOF in lower glycolysis and genes involved in the response to oxidative stress such as *prr1*, may explain the increased sensitivity to OS. However, some genes involved in OS response, autophagy and vesicle transport may contribute to cross-resistance to both stresses^97,98,112–114^ and indeed we observed three populations (G5, G9, and H9) with hits in those pathways that did not appear to have increased OS sensitivity and those populations retain WT-like expression level of the lower glycolysis pathway. This cross-resistance is also observed in other species like *K. lactis, S. cerevisiae*, and human cancer^115–118^. In *S. pombe*, the effect of the cross-resistance pathways was at times insufficient to offset the cost of adapting to HS by rewiring of the glycolytic flux toward fermentation and using it as the main source of biomass and energy. The fact that this trade-off has not been observed in the budding yeast may be explained by its historical exposure to multiple anaerobic environments and more optimal adaptations to HS^34^. In contrast, *S. pombe* and *S. japonicus,* for which lower glycolysis is less active compared to *S. cerevisiae*, tend to adapt to HS through LOF in lower glycolysis and upregulation of fermentation and upper glycolysis^32,35^. Furthermore, the fact that the response to HS mostly involved transcriptomic changes in genes that were not mutated in coding or flanking regulatory regions suggests that the adaptive transcriptomic changes observed across populations were driven by trans-regulatory rather than cis-regulatory effects. This also shows that the transcriptome, an intermediate level of organization between genotype and phenotype, is a complex trait that can exhibit evolutionary parallelism.

Despite the potential of these observations to improve evolutionary predictions, this study shows that multiple mechanisms are involved and that contingency can lead to diverse evolutionary outcomes, even within a single species and under the same conditions. One way that future research could better predict adaptive trajectories under stress conditions is by leveraging the presence of rapid transcriptomic changes in response to a new stress independently of mutations, e.g. expression plasticity. These responses could provide hints toward adaptive cellular changes that could later be cemented by mutations with similar effects, as is the case in bird species that explore high-altitude conditions^11,119^.

## METHODS

### Experimental evolution and sequencing

We evolved 15 *S. pombe* populations of the laboratory strain 972h-at 30 °C in wells of a lidded, flat-bottom 96-well plate containing 128 μl of YPD plus antibiotics (1% yeast extract, 2% peptone, 2% dextrose, 100 μg/mL ampicillin, 25 μg/mL tetracycline) for 10,000 generations, collecting and freezing samples at -80 °C every 70 generations. The remaining wells contained blank controls or populations of other yeast species and will be described elsewhere. This is the same procedure used to generate the compared *S. cerevisiae* dataset^3^. Because these conditions yield a doubling rate of 10 generations per day, we performed a 1/1024 serial dilution every day (1/2^10^). This corresponds to an effective population size of ∼6.00E04 and a bottleneck population size of ∼8.00E03 ^3,120^. We chose to perform whole-population whole-genome sequencing every 1,000 generations, and we were able to grow back cells from 93 out of these 100 samples after storage and sequenced these time points with Illumina NovaSeq paired-end sequencing (mean effective coverage ∼40X). Sequencing and DNA preparation was performed as in the *S. cerevisiae* dataset. Raw DNA sequences are available on NCBI: https://www.ncbi.nlm.nih.gov/bioproject/PRJNA1315931.

### Measuring fitness in the *S. pombe* populations

We grew the evolved populations, the WT, and a *S. cerevisiae* reference strain expressing GFP (YAN438: *MAT*α *his3*Δ1 *ura3*Δ0 *leu2*Δ0 *lys2*Δ0 *can1*::*RPL39pr-ymGFP*_*STE2pr*-*SpHIS5*_*STE3pr-LEU2*), for 24 hours. In contrast to our previous study^3^, this *S. cerevisiae* strain does not express a killer phenotype as it does not possess ScV-M1 killer-virus RNA. Next, because the green *S. cerevisiae* strain grew much faster than *S. pombe,* we prepared mixtures of the “dark” *S. pombe* populations and the green budding yeast at a 9:1 ratio and grew the mixed cultures for 24 hours. This ratio also reduces potential species interactions between *S. cerevisiae* and *S. pombe*. We then serially propagated each mixture for 3 days and used a Beckman-Coulter CytoFlex flow cytometer to measure the frequency of the “dark” *S. pombe* and the green *S. cerevisiae* strain daily. All particle counts were acquired at 30 μL/s rate for a maximum of 60 seconds or until 10,000 events were detected from a 96-well plate containing the mixtures. The data were analyzed using the CytoFlex acquisition and analysis software CytExpert. The “dark” *S. pombe* cells were distinguished from the green *S. cerevisiae* strain using FITC-A/PE-A gates. This process was replicated on a different day for biological replicates. Using the cell counts, we then calculated the frequency of each population in the mixture and inferred fitness based on the slope of a linear regression between the logarithm of the frequency ratio and the number of generations:

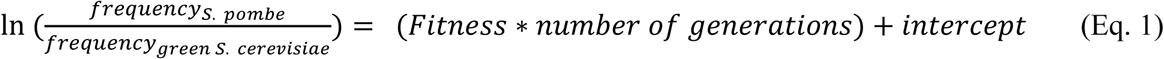

The final fitness was the average of the two biological replicates.

### Variant calling

We used Varscan2.4.6^47^ to call single-nucleotide variants (SNVs) and indels. Because contamination, sequencing errors and low-complexity regions can affect the variant calling, we applied some filters to the called variants (Table 2). These filters ensure that the calls are supported by reads, are time-consistent, and are not affected by possible errors in the reference genome, e.g. avoiding confusion between ancestral alleles that were not present in the reference genome (972h-) and fixed de novo mutations. Briefly, the filters used mapping and read base quality, allele frequency and absence of strand biases. In addition, because of the relatively sparse sequencing of our experiment (every 1000 generations), most genuine mutations will be fixed in cohorts or throughout the experiment, we also employed a filter for genetic hitchhiking and clonality.

**Table 2.**
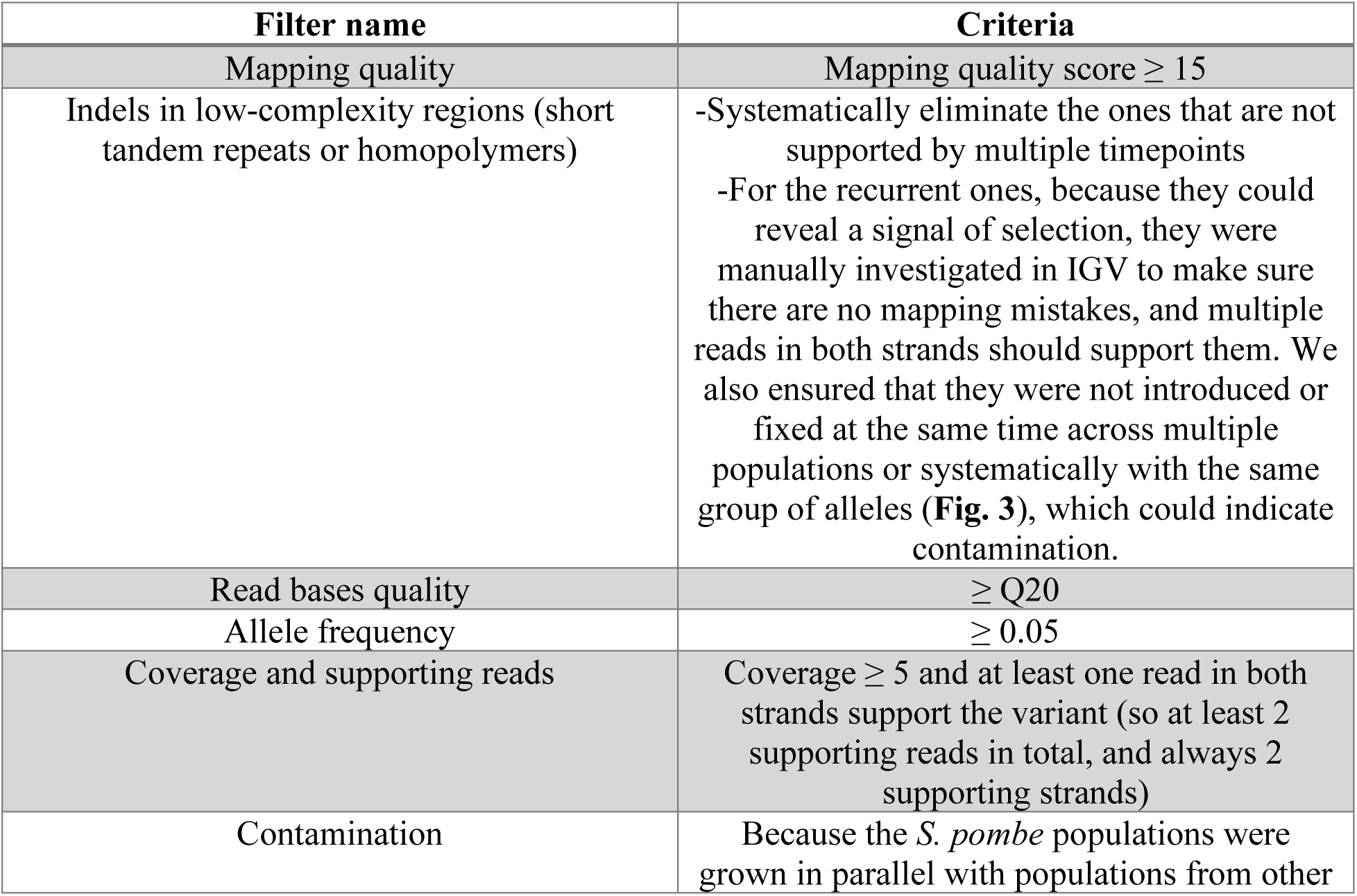
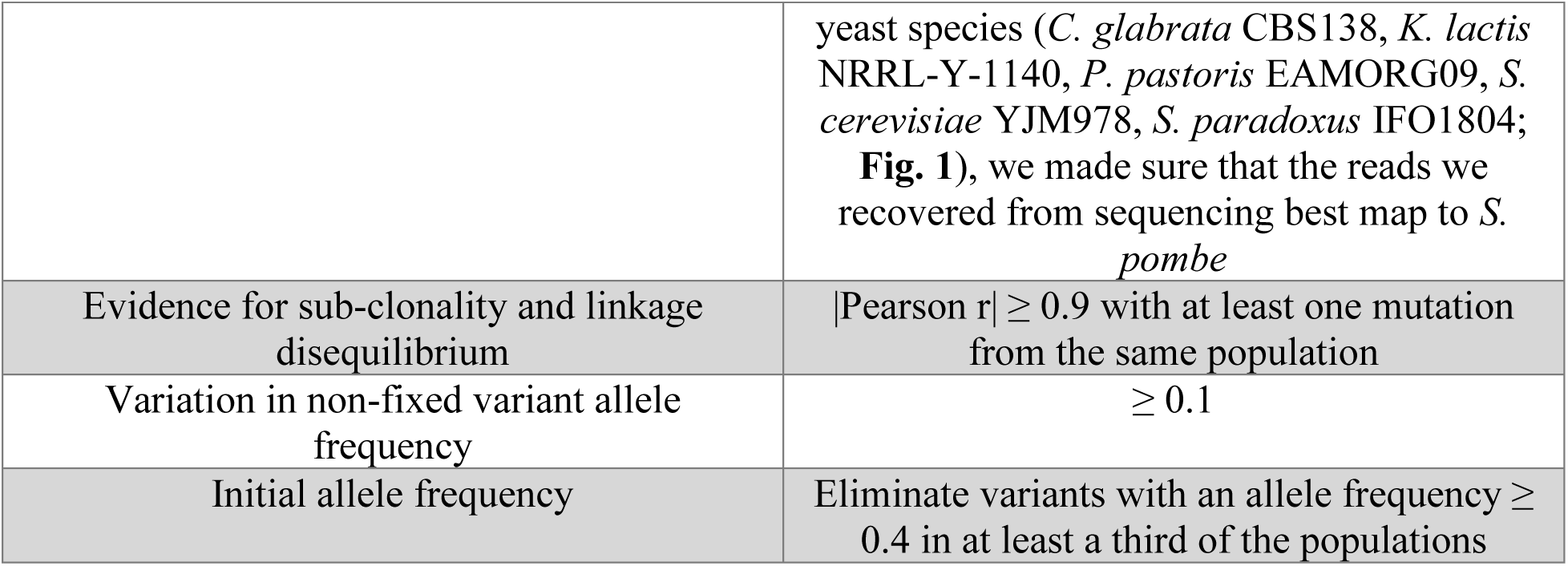
List of variant calling filters.

When applying these filters, we also found 10 cases where the same recurrent indel in low-complexity region was called in multiple populations. While these can be signatures of strong mutation bias or strong selection, they can also indicate evidence of cross-contamination in the evolution or DNA sequencing, as well as mis-indexed reads or read mapping issues. To rule these out, we applied several manual inspections. First, when mutations arise in a population and fix, they must remain fixed for the rest of the evolution. Second, if a recurrent mutation fixes in two populations, they must not substantially share other mutations within the samples and the rest of the time course (which would indicate cross-contamination). And third, other populations must not show strong evidence of these mutations (which would indicate some sequencing bias). While this does not rule out all possible issues with long-term evolution experiment and robotic passaging, or other artifacts of our variant calling pipeline, these mutations were still considered in our analyses as they may reveal mutational biases or other interesting phenomena (such as genetic-level nucleotide convergence). Despite these filters, three variants remain ambiguous (in *dep1, rpp1, pqr1*). A version of **Fig. 3** without these recurrent mutations is found in **Supplementary Fig. 7**.

We also found cases of mutations that appeared to remain at intermediate frequencies, one of which was recurrent (e.g. *ctf18*). This could indicate some form of frequency dependence or aneuploidy, but it can also be a signature of a read mapping artifact. To explore this, we manually inspected the mapped reads and found that these occurred in large polynucleotide tracts. As reads may not always span the whole tract, variant calling algorithms may fail to capture indels as fixed. We decided to leave the estimated frequencies as is.

Finally, complex mutations, such as events that may replace several adjacent codons simultaneously, may be due to alternative read mapping locations or due to complex short-tract recombination. We used the mapping quality filter to remove cases of alternative read mapping locations, and thus these complex mutations were left as is by the variant calling pipeline as they may be genuinely multiple mutations in the same region that get fixed. Nevertheless, this means that some categories of mutations may be inflated due to incorporation from a complex event.

The variant calling pipeline is available at https://github.com/arnaud00013/Experimental_evolution_S_pombe.

### Identifying signatures of selection from the substitution spectrum

We defined substitutions as variants with an allele frequency >= 75%. Then, we compared the substitution spectrum of each population with the one expected after 10,000 generations of evolution under a neutral null model. The number of fixed variants in each mutation category is calculated based on the species’ evolution rate, genome size, genome architecture (e.g. ratio of coding to non-coding regions in the genome), and the proportion of single-nucleotide mutations derived from the genetic code and associated with a specific category of mutations. This allowed to derive 2 equations:

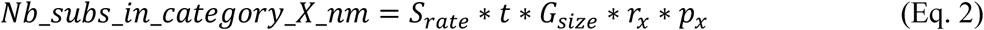

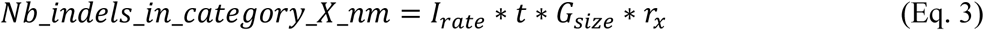

where 𝑁𝑏_𝑠𝑢𝑏𝑠_𝑖𝑛_𝑐𝑎𝑡𝑒𝑔𝑜𝑟𝑦_𝑋_𝑛𝑚 is the expected number of substitutions in a mutation category X assuming a neutral model of evolution, 𝑁𝑏_𝑖𝑛𝑑𝑒𝑙𝑠_𝑖𝑛_𝑐𝑎𝑡𝑒𝑔𝑜𝑟𝑦_𝑋_𝑛𝑚 is the expected number of indels of type X assuming a neutral model of evolution, 𝑆_𝑟𝑎𝑡𝑒_ is the species substitution rate, 𝐼_𝑟𝑎𝑡𝑒_ is the species indel rate, t is the number of generations observed since the start of the experiment, 𝐺_𝑠𝑖𝑧𝑒_ is the species genome size, 𝑟_𝑥_ is the proportion of the genome covered by protein-coding or non-coding sites, depending on the mutation category X and 𝑝_𝑥_ is the proportion of single-nucleotide mutations derived from the genetic code that can be classified as mutations from category X.

Next, we sought signatures of parallelism and positive selection in protein-coding genes among the fixed variants. We defined a “hit” as a fixed variant that is not synonymous (missense, nonsense, or indel), and a multi-hit gene as a gene with hits in multiple populations. (in at least 2 out of 10 populations). We then compared the number of population hits (PH) in the set of multi-hit genes to the PH expected under a null model, assuming that all genes evolve and fix variants at the same rate. The expected PH was determined based on a number drawn from a Poisson distribution where the argument λ (average number of events per interval) is the product of S. pombe’s substitution rate, the gene length, the generation time, the number of populations, and the proportion of fixed variants that are not synonymous as determined in equations 2 and 3.

### Evaluating and comparing the sensitivity to hydrogen peroxide

We grew the 10 evolved *S. pombe* populations and the WT ancestor in the rows of a 96-well plate at 30°C for 36 hours. The media was YPD + hydrogen peroxide with H_2_O_2_ concentrations of 0 mM, 0.5 mM, 1 mM, 1.5 mM, 2 mM, 2.5 mM, 3 mM and 3.5 mM. Each row of the 96-well plate corresponded to a different concentration of H_2_O_2_. The growth of each population was estimated using the OD600 time series measured from a BioTek Synergy HTX plate reader. For each time series, the area under the curve (AUC) was measured using the R package package growthcurver^121^. Using the AUC as a proxy for growth, we then compared the AUC of each population at different H_2_O_2_ concentrations to the AUC obtained when H_2_O_2_ is absent from the media. Next, we inferred the degree of inhibition with:

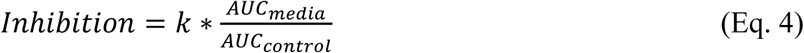

where 𝑘 is a normalizing factor that makes sure the inhibition metric maximum is 1, AUC is the area under the growth curve, 𝐴𝑈𝐶_𝑚𝑒𝑑𝑖𝑎_ represents the growth at a certain concentration of hydrogen peroxide and 𝐴𝑈𝐶_𝑐𝑜𝑛𝑡𝑟𝑜𝑙_ represents the growth in a YPD media without hydrogen peroxide. This expression is negatively correlated to the AUC ratio and ranges from 0 to 1. Using the Dose-Response Analysis R package drc^122^, we then fitted a log-logistic model to this metric and H_2_O_2_ concentrations to estimate the IC50, which recapitulates the level of sensitivity of a population to hydrogen peroxide. Data in the manuscript is a representative result of the less noisy biological replicate as determined by curve fit. Additionally, we found that the hydrogen peroxide concentration tended to vary due to instability of the chemical, which would at times shift the IC50, without changing the interpretation of the experiment. Finally, we performed a cluster analysis on the populations based on their distribution of AUC media-to-control ratio at different concentrations. We used the hclust function from the package stats v4.1.2^123^ to perform the clustering and the composition of each cluster was identical no matter the distance metric tested (Gower, Manhattan, Euclidean, Bray-Curtis), which we obtained with the function vegdist from the package vegan 2.7-2^124^. Growth curve figures are also shown in **Supplementary Fig. 8**.

### RNA extraction, sequencing and transcriptomic analysis

We extracted in two biological replicates the RNA of the evolved populations using an in-house protocol that works well with *S. cerevisiae*. Briefly, cells were grown to mid-log phase in YPD by diluted overnight saturated cultures and growing for 4-6 hours and lysed using 100 μl of 5 mg/mL Zymolyase + 10 mM DTT, incubated at 37 °Celsius for 5 minutes. The weakened cells were then incubated with 100 μl of 2% SDS for a final SDS concentration of 1%. 200 μl of lysis buffer was added (4.5 M Guanidine thiocyanate, 10 mM EDTA) and mixed gently by inverting. Lysate clarification indicated complete lysis. The SDS was then precipitated using 200 μl of 3M Potassium acetate at pH 5.

After centrifugation, the lysate was passed through a minipreparation silica column to bind DNA. Under these conditions, RNA does not bind to the silica columns. Therefore, we collected the flowthrough. We then mixed it with an equal volume of pure isopropanol and passed it through another minipreparation silica column (which will now bind RNA). The column was then washed with 400 μl of wash buffer (10% Guanidine thiocyanate, 25% Isopropanol, and 10 mM EDTA), and then twice with 600 μl of 80% Ethanol. After drying the column, the RNA was eluted with 50 μl of 10 mM Tris pH 8.5. RNA integrity was verified on an agarose gel. One population (H12) failed to get sufficiently high-quality RNA for sequencing.

We sequenced the successfully extracted RNA using NEBNext Ultra II on an Illumina NovaSeq X paired-end sequencing lane. This yielded 10.6E06 150-bp paired-end reads per sample on average (σ=1.32E06). Raw RNA-sequencing data are available on NCBI: https://www.ncbi.nlm.nih.gov/bioproject/PRJNA1315931. We pre-processed the reads with fastp (version 0.23.1) with the arguments --trim_poly_x --trim_poly_g --cut_front --cut_front_mean_quality 20 --cut_tail --cut_tail_mean_quality 20 --detect_adapter_for_pe^125^. Next, we mapped the reads to the reference (972h-) using samtools v1.17, removed duplicates with picard v2.26.3 and counted the number of reads per gene using featureCounts from subread v2.0.6^126–128^, and the counts were converted to FPKM. We then performed PCA analysis of the transcriptome and found two main clusters (PC1 and PC2 explained about 75% of the transcriptomic variance), one which contained the WT and the other contained only populations that were more sensitive to oxidative stress than WT. One population that was as sensitive as WT clustered independently, and two populations that clustered with the WT were more sensitive to oxidative stress. Upon analysis of their gene hits, we found that they have loss-of-function mutation in genes that relate to oxidative stress resistance.

To determine which genes and their expression differences may be responsible for the increased oxidative stress sensitivity of most populations, we used the DESeq2 package to detect differentially expressed genes between the two clusters (i.e. populations that are as sensitive and more sensitive than WT)^129^. Gene functional categories and GO annotations were obtained from Pombase data^40^. The RNA-seq analysis pipeline is available at https://github.com/arnaud00013/Experimental_evolution_S_pombe.

