## Supplemental Figures S1-S8 for "Parallel but distinct adaptive routes in the budding and fission yeasts after 10,000 generations of experimental evolution"

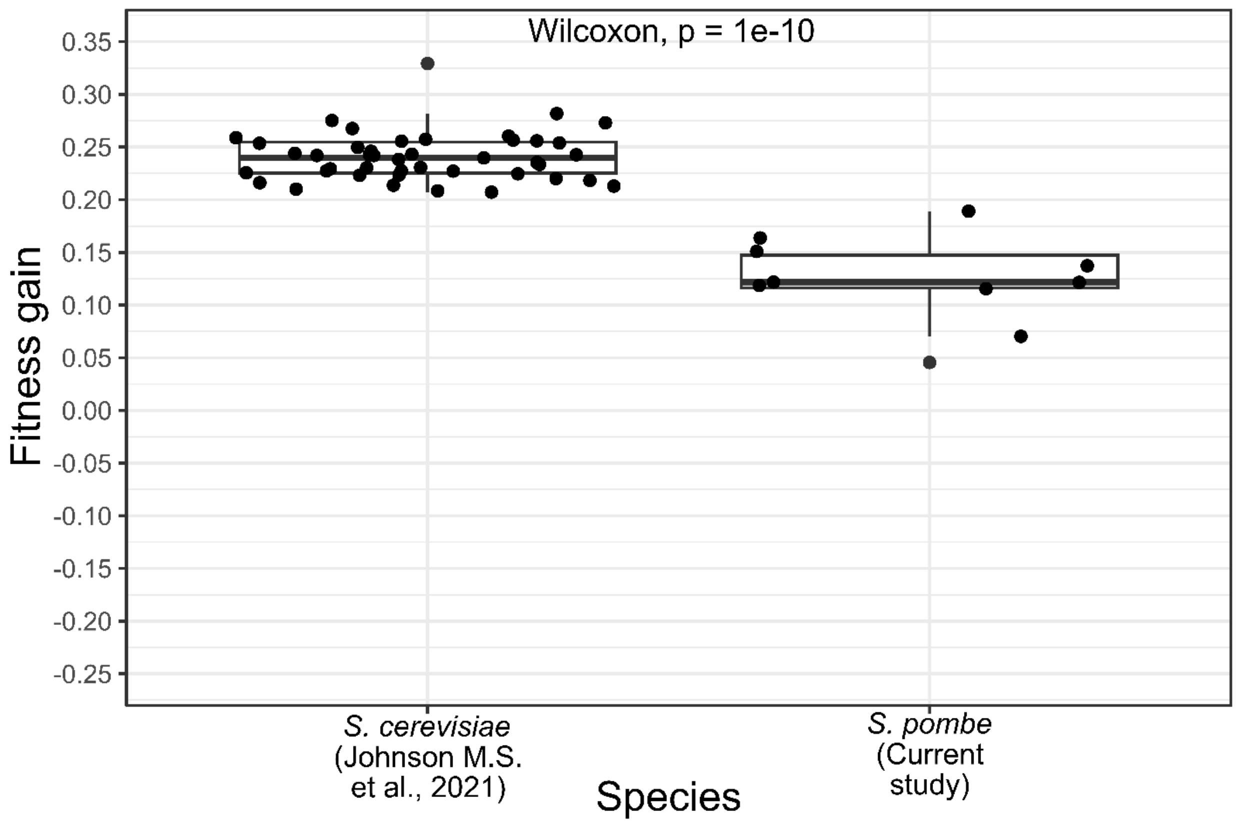


**Supplementary Figure 1: Fitness gain measured during experimental evolution of *S. cerevisiae* and *S. pombe***. The fitness gain is the difference between the fitness at the final time point (10,000 generations) and the ancestral fitness (time 0). The two-sided p-value is obtained from performing a Wilcoxon rank sum test (Mann-Whitney U test).

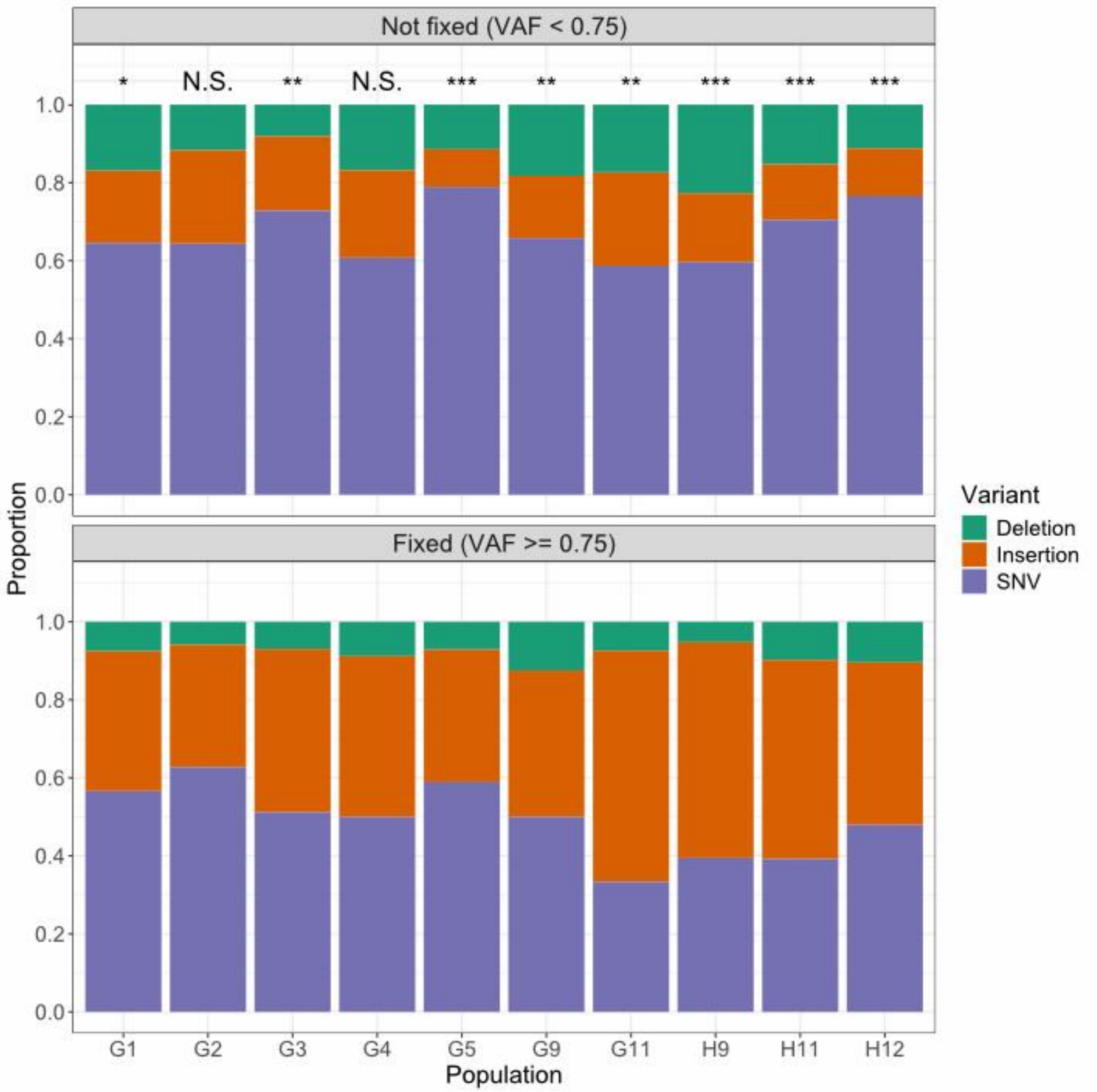


**Supplementary Figure 2: *S. pombe* fixation bias**. The fixation bias is defined as the change of frequencies between the fixed and non-fixed variants. The two-sided p-values of a Chi-Squared test are encoded as follows: *** from 0 to 0.001 exclusively, ** from 0.001 to 0.01 exclusively, * from 0.01 to 0.05 exclusively, "." from 0.05 to 0.1 exclusively and N.S. otherwise, which is the acronym of "non-significant".

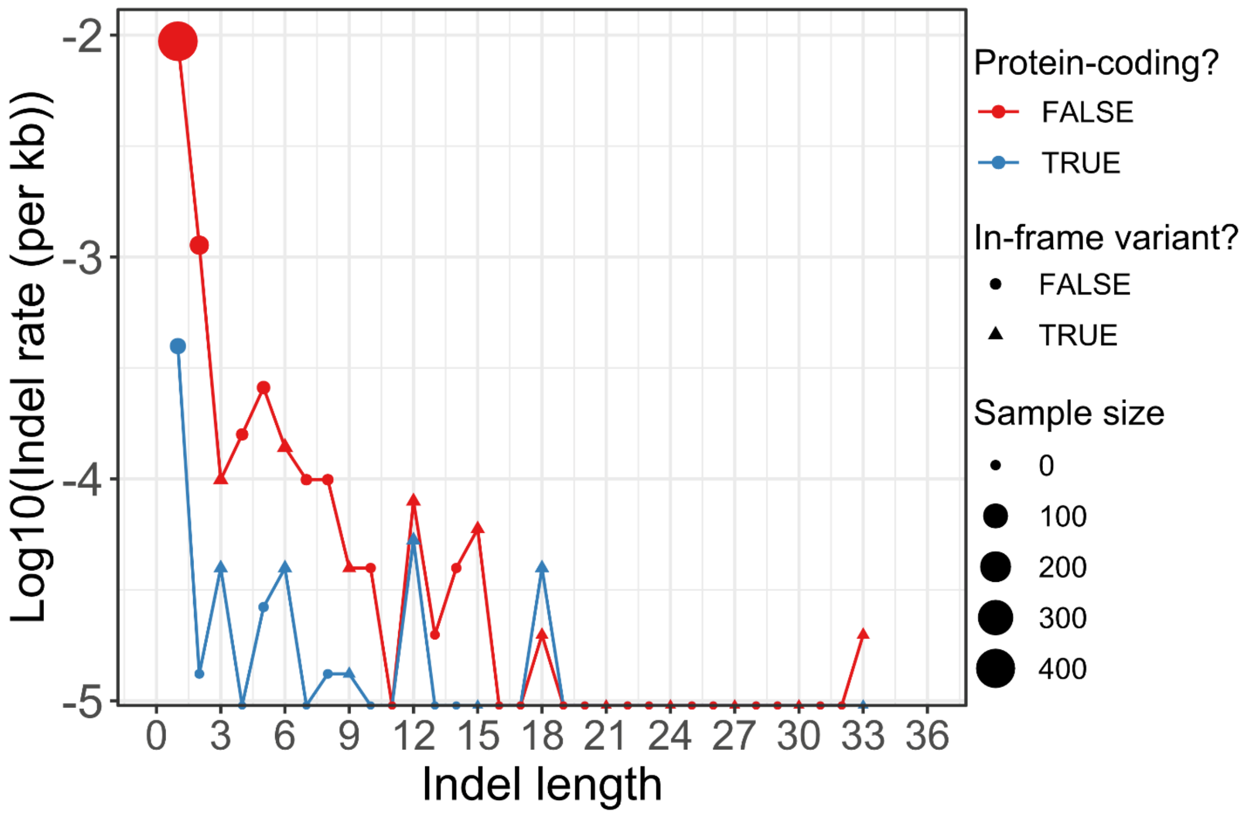


**Supplementary Figure 3: Indel rate for coding vs non-coding indels and in-frame vs frameshift indels**. We defined the indel rate as the number of indels per kilobase (Kb) and plotted it on a log10 scale on the y-axis. The dot size represents the size of each indel set.

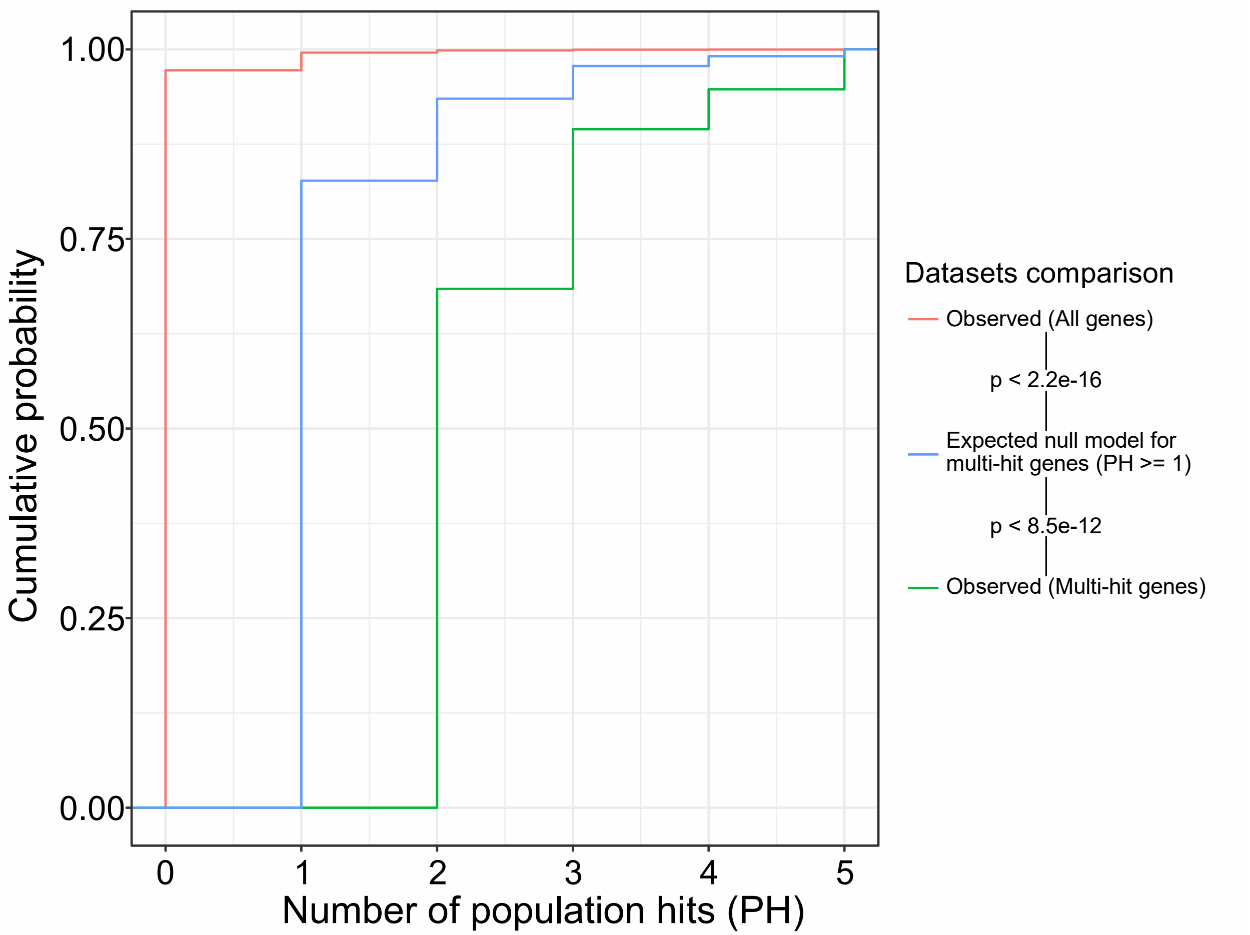


**Supplementary Figure 4: The expected vs observed cumulative distribution of population hits across the 10 *S. pombe* populations**. We defined population hits (PH) as the number of populations in which a gene has a hit. The null model assumes that all genes evolve and fix variants at the same rate, so the expected PH is determined by this assumption and follows a Poisson distribution (**Methods**). The one-sided p-values were obtained using a Kolmogorov-Smirnov test (alternative: cumulative distribution function 1 is "higher" than function 2, which means that there are higher values in function 2).

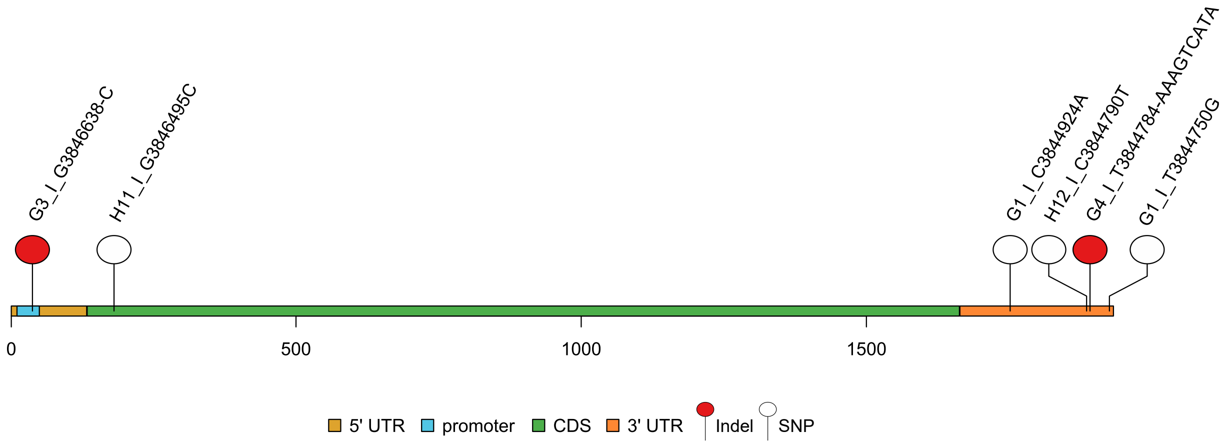


**Supplementary Figure 5: *pyk1* mutations across the 10 evolved *S. pombe* populations**. The positions in the pyruvate kinase gene *pyk1* are illustrated on the x-axis while the coding and non-coding regions have distinct colors. The mutations are red when they represent indels and white when they represent single-nucleotide polymorphisms. The format of the variant name is "population_chromosome_reference allele_position in the chromosome_new allele". For insertions, the alternative allele starts with the character "+" while an alternative allele starting with the character "-" represents a deletion.

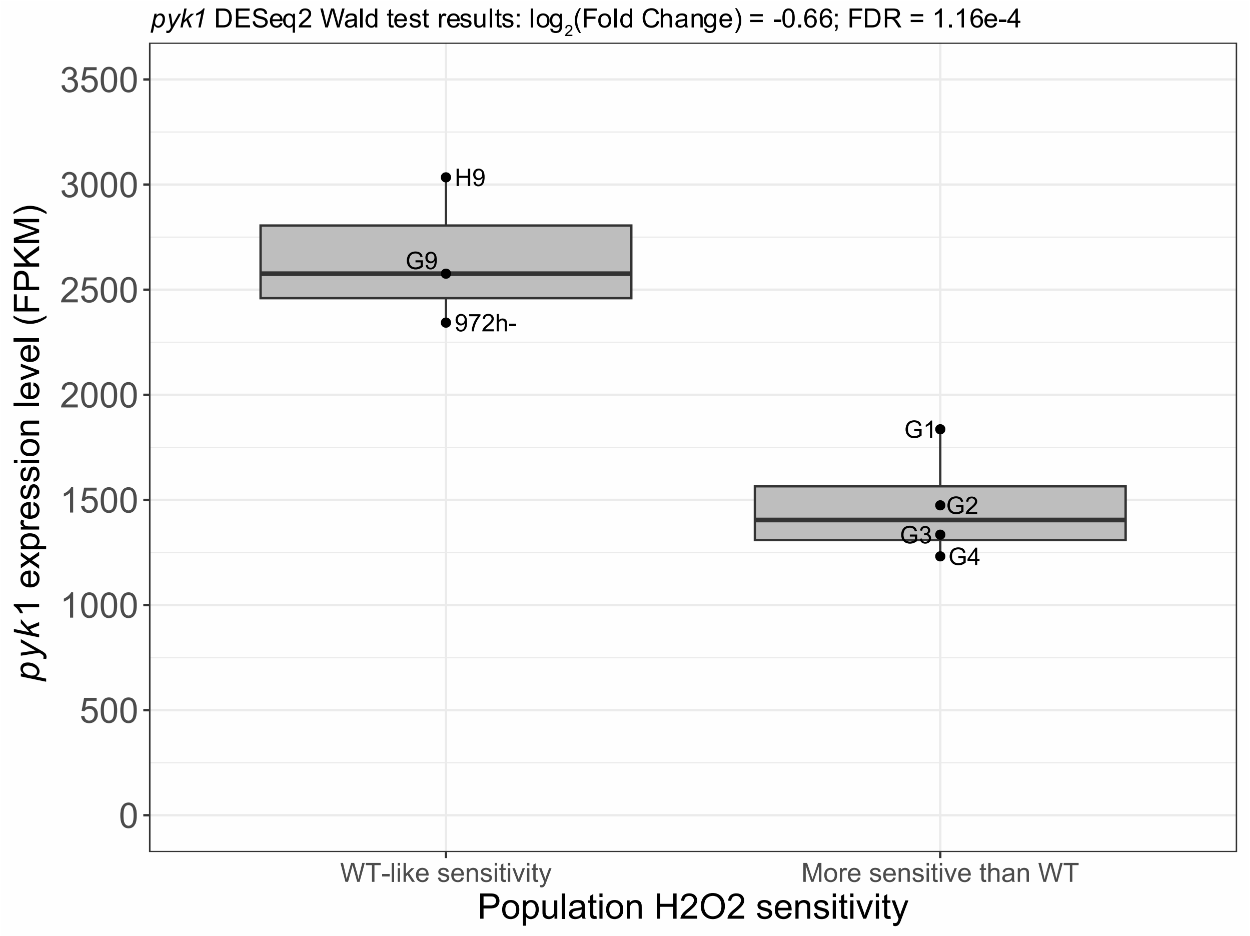


**Supplementary Figure 6: *pyk1* expression level in WT-like populations compared to populations that are more sensitive to OS**. Gene expression levels are obtained from read mapping (**Methods**) using the number of Fragments Per Kilobase of transcript per Million mapped reads (FPKM) metric. We defined WT-like populations as evolved populations with transcriptomic profiles and OS sensitivities similar to WT. Populations that are more sensitive to OS than WT but that have a similar overall transcriptomic profile compared to the WT have been excluded from this analysis as their sensitivity to OS could be explained by LOF hits in protein-coding genes independently of expression (**Methods**).

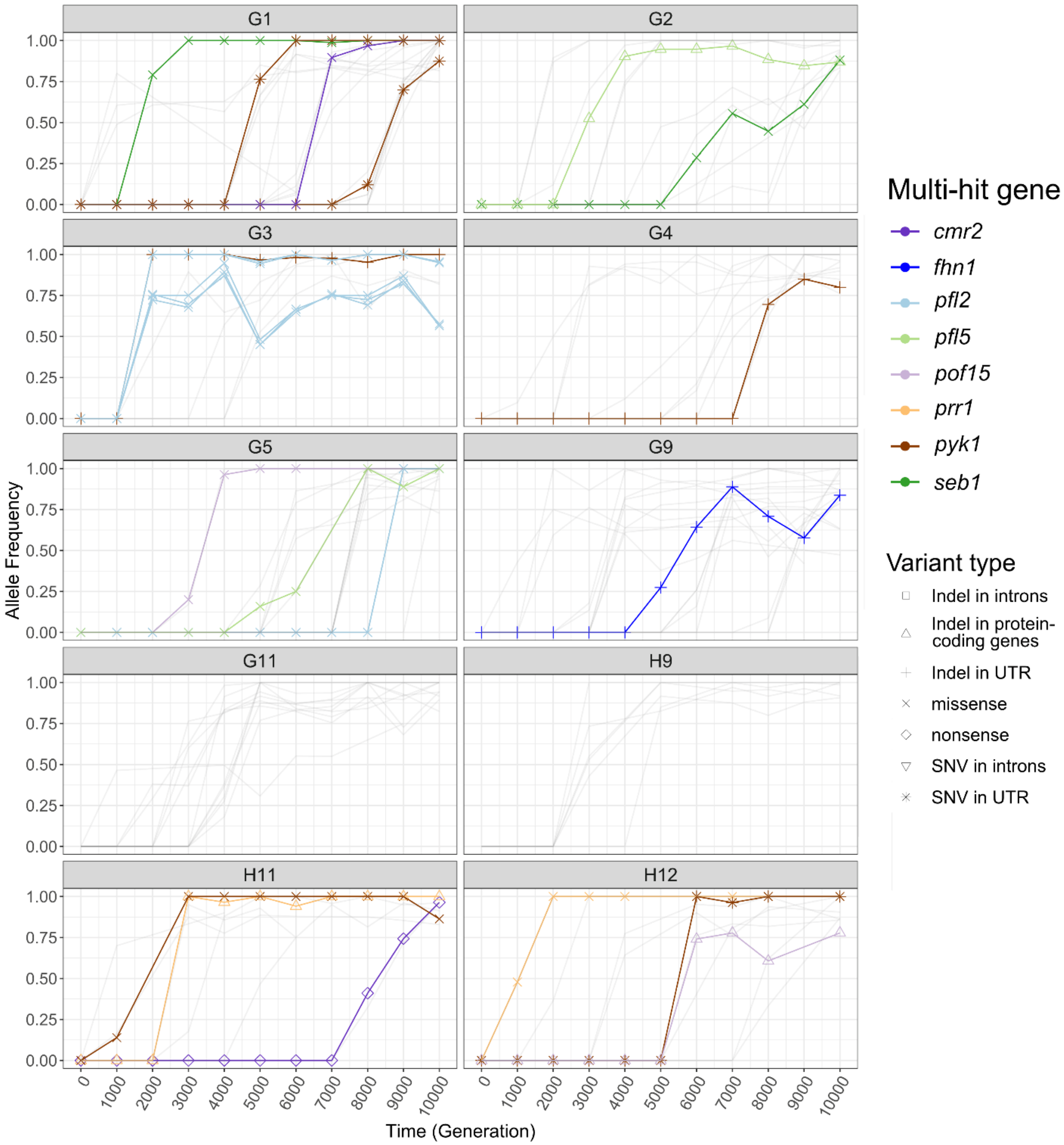


**Supplementary Figure 7: Time series of the hits' allele frequencies in evolved *S. pombe* populations and multi-hit genes without recurrent mutations**. Time series of the hits' allele frequencies in the *S. pombe* populations without recurrent mutations. We defined a "hit" as a fixed genic variant that is not synonymous (missense, nonsense and indels) and a multi-hit gene as a gene having hits in multiple populations. In each population, variants that were never fixed or that are not located in multi-hit genes are represented by more transparent lines. In the legend, PH represents the number of populations in which the gene is hit.

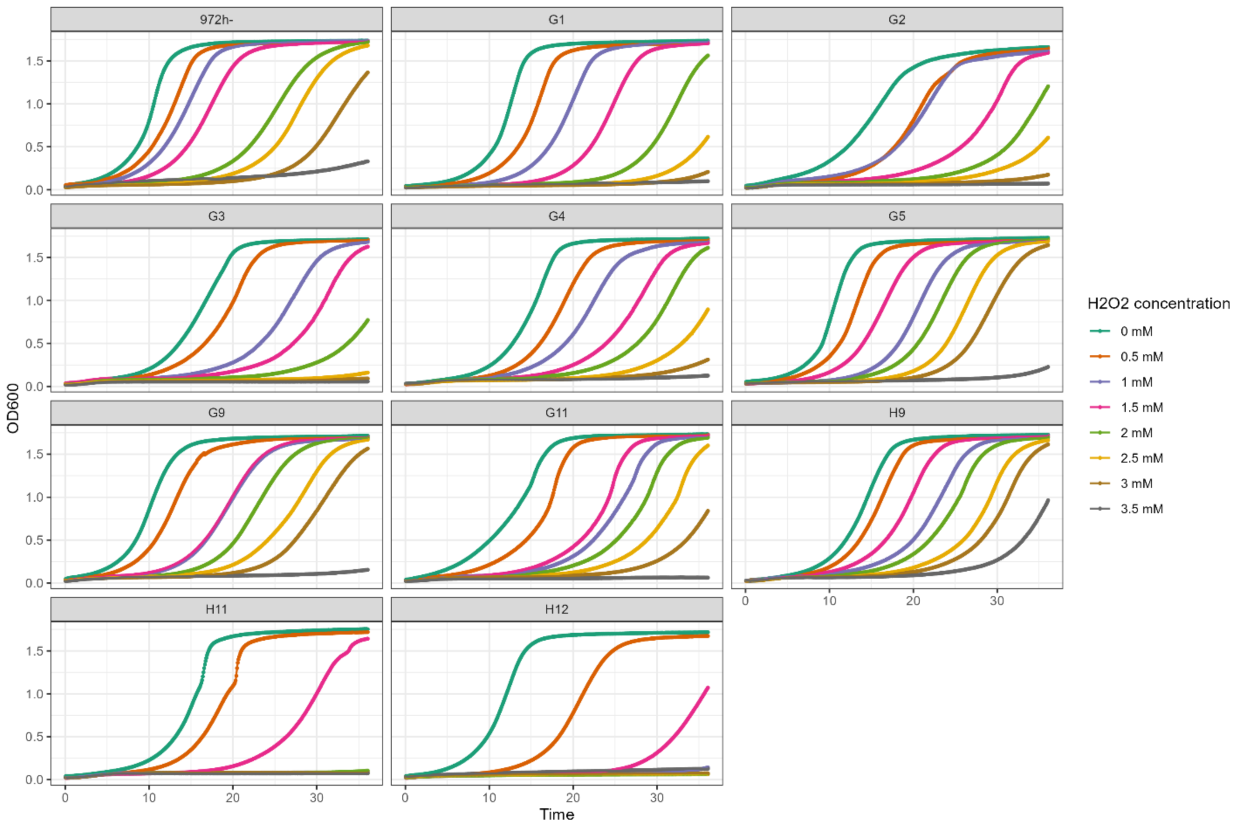


**Supplementary Figure 8: Growth curves of evolved populations at different hydrogen peroxide concentrations**. *S. pombe* evolved populations and ancestor were grown at different hydrogen peroxide concentrations for 36 hours to determine their resistance to oxidative stress.
